# Structural dynamics of the functional nonameric Type III translocase export gate

**DOI:** 10.1101/2020.11.20.391094

**Authors:** Biao Yuan, Athina G. Portaliou, Rinky Parakra, Jochem H. Smit, Jiri Wald, Yichen Li, Bindu Srinivasu, Maria S. Loos, Harveer Singh Dhupar, Dirk Fahrenkamp, Charalampos G. Kalodimos, Franck Duong van Hoa, Thorben Cordes, Spyridoula Karamanou, Thomas C. Marlovits, Anastassios Economou

**Author notes:** For correspondence: Anastassios Economou Thomas C. Marlovits. **Authors e-mail addresses: Biao Yuan:**, **Athina G. Portaliou:**, **Rinky Parakra:**, **Jochem H. Smit:**, **Jiri Wald:**, **Yichen Li:**, **Bindu Srinivasu:**, **Maria S. Loos:**, **Harveer Singh Dhupar:**, **Dirk Fahrenkamp:**, **Charalampos G. Kalodimos:**, **Franck Duong van Hoa:**, **Thorben Cordes:**, **Spyridoula Karamanou:**, **Thomas C. Marlovits:**, **Anastassios Economou:**.

## Abstract

Type III protein secretion is widespread in Gram-negative pathogens. It comprises the injectisome with a surface-exposed needle and an inner membrane translocase. The translocase contains the SctRSTU export channel enveloped by the export gate subunit SctV that binds chaperone/exported clients and forms a putative ante- chamber. We probed the assembly, function, structure and dynamics of SctV from enteropathogenic *E.coli* (EPEC). In both EPEC and *E.coli* lab strains, SctV forms peripheral oligomeric clusters that are detergent-extracted as homo-nonamers. Membrane-embedded SctV_9_ is necessary and sufficient to act as a receptor for different chaperone/exported protein pairs with distinct C-domain binding sites that are essential for secretion. Negative staining electron microscopy revealed that peptidisc-reconstituted His-SctV_9_ forms a tripartite particle of ∼22 nm with a N-terminal domain connected by a short linker to a C-domain ring structure with a ∼5 nm-wide inner opening. The isolated C-domain ring was resolved with cryo-EM at 3.1 Å and structurally compared to other SctV homologues. Its four sub-domains undergo a three-stage “pinching” motion. Hydrogen-deuterium exchange mass spectrometry revealed this to involve dynamic and rigid hinges and a hyper-flexible sub-domain that flips out of the ring periphery and binds chaperones on and between adjacent protomers. These motions are coincident with pore surface and ring entry mouth local conformational changes that are also modulated by the ATPase inner stalk. We propose a model that the intrinsic dynamics of the SctV protomer are modulated by chaperones and the ATPase and could affect allosterically the other subunits of the nonameric ring during secretion.

## INTRODUCTION

The type III protein secretion system (T3SS) is widely used by many Gram-negative pathogenic or symbiotic bacteria to inject proteins directly into the eukaryotic host cytoplasm (1, 2). T3SS comprises the “injectisome”, a nano-syringe bridging the bacterial and eukaryotic cytoplasm. Injectisomes comprise four parts (Fig. 1A): a ∼32-36 nm-long trans- envelope “basal body” composed of stacked rings; an inner membrane-embedded “export apparatus”, located at the basal body base; a circular cytoplasmic sorting platform, including the ATPase complex, that protrudes ∼28 nm intro the cytoplasm and has a diameter of ∼32 nm and finally, a filamentous needle lying at the top of the injectisome, that protrudes from within the basal body to the extracellular milieu, and carries a “translocon” at its tip (2–5). Translocons form pores in host plasma membranes through which effectors are delivered (6).

**Fig. 1.**
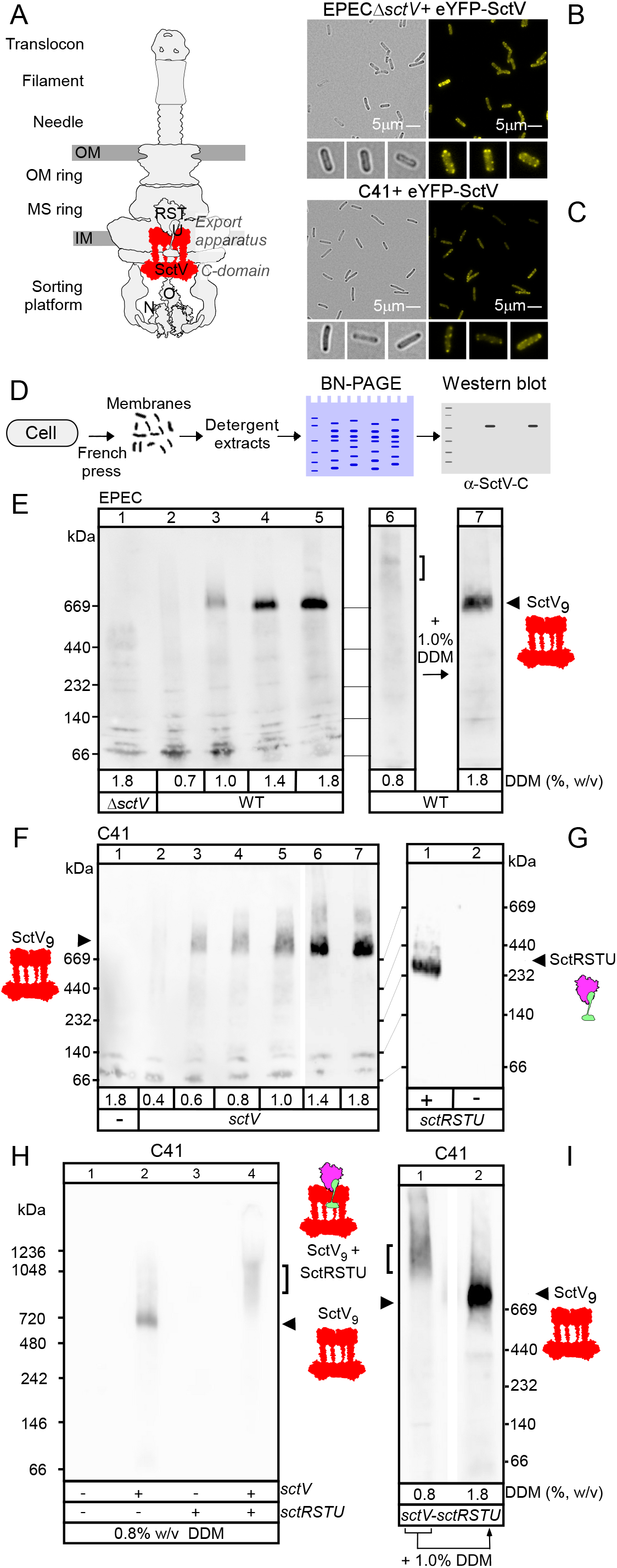
Subcellular localization of SctV and characterization of SctV nonamers. **A.** Cartoon representation of the EPEC injectisome. SctV_9_ is shown in red. **B** and **C.** eYFP-SctV forms distinct foci in EPECΔ*sctV* (**B**) and C41(**C**) cell peripheral. Scale bar: 5 μm; *eYFP-sctV* expression was induced with AHT: 20 ng/mL Representative images from YFP and brightfield channel are shown; *n=3* biological replicates. **D:** Pipeline for SctV membrane complexes analysis by BN-PAGE. Following French press, lysed cells were removed and membrane fractions were harvested by ultra-centrifugation. Non-ionic detergents DDM was used to extract membrane proteins for further analysis by BN-PAGE and immuno-staining with α-SctV-C. **E.** BN-PAGE analysis of EPEC-derived, DDM-solubilized membrane complexes containing SctV. Membrane proteins were extracted with different concentrations of DDM, analyzed by BN-PAGE and α-SctV-C immuno-staining (lanes 1-6). ∼1 MDa species containing SctV component was seen when extracted with 0.8% DDM (lane 6, bracket). A predominant ∼700kDa species also appeared when extra DDM was added into 0.8% DDM sample (lane 7) migrating on the gel with the similar mass size as extractions with ∼1.0% DDM (lanes 3- 5). Representative images are shown; *n=3* **F.** As in E, DDM-solubilized membrane complexes derived from C41/ *sctV* cells. Membrane proteins were extracted with different concentrations of DDM, analyzed by BN-PAGE and α-SctV-C immuno-staining. Representative images are shown; *n=3* **G** SctRSTU complex formation in C41 cells. BN-PAGE analysis of C41 or C41/*sctRSTU*, Digitonin-solubilized membrane complexes and α-SctU-C immuno-staining. A representative image is shown; *n=3* **H-I.** SctRSTU-V complex formation in C41 cells. Complexes derived from C41cells carrying compatible plasmids with *sctV* and *sctRSTU* (expressed as indicated) extracted with 0.8% DDM **(H)** or further treated with 1% DDM **(I)** and analyzed by BN-PAGE and α-SctV-C immuno-staining. Representative images are shown; *n=3*

The unifying nomenclature Sct (<u>S</u>ecretion and <u>c</u>ellular translocation) is used for the conserved components of the pathogenic/symbiotic injectisomes across bacteria (7, 8)(www.stepdb.eu). Species/genus suffixes in subscript are added here to denote Sct components from other bacteria (e.g. *_Sf_*Sct for *Shigella flexneri*).

The export apparatus located centrally inside the injectisome is thought to be the translocase that mediates chaperone/exported client targeting, switching from one client class to another, client engagement and translocation through the secretion tunnel (2, 9, 10). Five components (SctRSTU and V) of the translocase contain many hydrophobic transmembrane-like domains, which challenge structural characterization and functional definition. This is further complicated by the peripheral nature of the association of SctV with SctRST seen from cryo-ET analysis (3, 5, 11). SctV and U also contain large C-terminal cytoplasmic domains. The SctV export gate is a nonamer and its C-domain forms a large cytoplasmic ring with a ∼5 nm vestibule connected to the N-terminal membrane part with narrow stalks (12–15)(Fig. 1A). The isolated cytoplasmic domain of *Shigella flexneri _Sf_*SctV- C crystalized as a ring structure with an external diameter of 11-17 nm and a height from the membrane surface of ∼5.5 nm (12, 16, 17). Similarly, the flagellar SctV(FlhA)-C forms rings with an external diameter of 10.1 ± 0.6 nm and a height of 5.1 ± 0.4 nm (18–21). The C-terminal domain of _EPEC_SctV has been structurally resolved at 4.6 Å resolution using cryo- EM revealing that nonameric rings form via electrostatic interactions between subunits (14). The nonameric *_St_*SctV of *Salmonella typhimurium* has been determined using a high- throughput cryo-ET pipeline (16, 22). Such a cryo-ET model still lacks sufficient resolution to determine function and the assembly of its transmembrane channel formed around other subunits of the export apparatus by 72 transmembrane helices. Notably, the ATPase is located at a significant distance from SctV [∼5 nm; (3, 5, 11)], but may connect to the SctV cytoplasmic ring indirectly through the elongated SctO inner stalk (23–27). Additional peripheral bridges formed by other components of the cytoplasmic sorting platform keep it in place juxtaposed to SctV even when the stalk is missing (3). The dynamics of ATPase/stalk/SctV C-domain during the translocation cycle remain unknown.

The monomeric cytoplasmic domains of SctV/FlhAs(SctV-C/FlhA-C) expressed as individual polypeptides interact with chaperone/exported protein complexes in solution (19, 28, 29) with a low dissociation constant (*K_d_)* of ∼20 μM (29). Chaperone/exported protein complexes also interact with the hexameric ATPase of the system in NMR-identified complexes (30) and with cytoplasmic platform components in pull-down experiments (26, 31, 32). Although the order of chaperone/exported protein interactions with machinery components during secretion remains debatable, such interactions presumably attract exported substrates to SctV (9, 29). Then exported substrates dissociate from chaperones, diffuse to the SctV vestibule and to the inner membrane SctRSTU for export, via a putative channel running longitudinally through SctRST (10, 13, 33, 34). Recently, the first structure of an active export apparatus engaged with an exported protein in two functional states was reported, revealing the complete 800 Å-long secretion conduit and unravelling its critical role in T3 secretion (13). However, export protein binding and trapping steps are poorly understood and the precise order and roles of the translocase components are unknown.

cryo-ET revealed that within the assembled injectisome, *_St_*SctRST sits on top of SctV in an inverted cone formation that is largely periplasmically exposed and only partially buried in the inner membrane plane (3, 13, 16). It is kept in this periplasmically protruding state apparently through contacts with the SctJ lipoprotein and lipids (3). A similar arrangement was seen in the flagellar T3S (33). The intricate formation of the injectisome rings would imply possible architectural dependence of one sub-structure on the other and perhaps suggest a temporal order of addition of each element. Nevertheless, deletion of *sctV* does not prevent the correct assembly of the remaining injectisome including that of the cytoplasmic sorting platform, the SctRST inverted cone, that should be internal to SctV ring, and the SctD that surrounds SctV externally (16, 22). Moreover, the injectisome can be purified with non-ionic detergents in stable structures in the apparent absence of SctV (13, 15, 35, 36). These data suggested that assembly of SctV may retain a significant autonomy from that of the other architectural components.

Using the Gram-negative pathogen enteropathogenic *Escherichia coli* (EPEC), a serious diarrhea threat to children in developing countries (37), we recently reconstituted T3SS substrate membrane targeting process *in vitro* using inverted inner membrane vesicles (IMVs) harboring EPEC injectisomes. We revealed a novel mechanism that governs both membrane targeting of exported clients and switching from translocator (middle client) to effector (late client) export (9). We now focus on the assembly, structure, structural dynamics and receptor function of the export gate. We report that nonamers of *_EPEC_*SctV and of its C-domain self-assemble *in vivo* in the absence of any other T3S components, with the C-domain having a major contribution to nonamerization. Membrane-embedded SctV_9_ is necessary and sufficient to act as a functional receptor for chaperone/exported protein complexes in the absence of ongoing secretion and/or other injectisome components. Specific patches on its C-domain are important for chaperone/ exported protein complexes to bind as demonstrated using immobilized peptide arrays and confirmed with mutational analysis. The structure of the isolated C-domain polypeptide was resolved at 3.1 Å resolution by cryo-EM. Negative staining EM analysis indicates that peptidisc-reconstituted full-length His-SctV_9_ is a two-domain particle connected by an equatorial constriction. Both peptidisc-reconstituted His-SctV_9_ and cryo-EM-resolved C- domains showed nonameric rings with an internal pore of a ∼5 nm diameter. Important contacts between the protomers are essential for C-domain nonamerization. The C-domain architecture revealed four sub-domains connected by linkers and compared to other homologue structures suggest that they undergo an “open” to “closed” “pinching motion. Hydrogen-deuterium exchange mass spectrometry revealed extensive local intrinsic dynamics in parts of the structure that would allow rigid body motions and rotations including those of the hyper-flexible sub-domain 2 (SD2) that is mainly responsible for the pinching motion. SD2 is a non-conserved structure that overlooks a deep groove that separates two adjacent protomers and forms a highly negatively charged ridge on either side of the groove. This landscape defines a wide, multi-valent chaperone trap with multiple binding patches that are essential for secretion. Direct contacts of the ATPase complex inner stalk SctO would modulate both local and domain dynamics.

These findings lay the foundations for mechanistic dissection of how structural dynamic states are coupled to T3S translocase-mediated secretion.

## RESULTS

### SctV forms multimeric clusters *in vivo*

To analyze SctV membrane assembly *in vivo*, we fused SctV to the C-terminus of eYFP and placed it under tetracycline promoter-control (anhydrotetracycline; AHT)(9). EPECΔ*sctV,* a non-polar deletion strain, is similarly complemented *in vivo* by vector-borne *sctV* (Fig. S1A, lane 3), *eyfp-sctV* (lanes 5-6) and *his-sctV* (Fig S1B, lanes 6-8). eYFP-SctV assembles in distinct focal clusters in the cellular periphery detected by live-cell fluorescence microscopy (Fig. 1B), while eYFP alone shows widespread diffusion with no clusters (Fig. S1C).

To test if eYFP-SctV clustered assembly *in vivo* requires any other T3S components or if it self-assembles, we imaged the *E.coli* lab strain C41, that is devoid of an injectisome, carrying *eyfp-sctV*. C41/*eyfp-sctV* cells synthesized a similar amount of eYFP-SctV as EPECΔ*sctV/eyfp-sctV* (Fig. S2A) that became visible after ∼20 min (Fig. 1C; Fig. S2B). This presumably also encompasses the maturation time for the fluorophore (38). eYFP-SctV synthesized in C41, as in EPECΔ*sctV* (Fig. 1B), also showed a distinct peripheral punctate staining (Fig. 1C). This indicated that the apparent eYFP-SctV clustering *in vivo* is an inherent property of SctV.

The punctate staining pattern of eYFP-SctV was characteristic and distinct from that of other fluorescent fusion protein markers that showed either diffuse staining all over the cytoplasm (eYFP) or peripheral staining but of similar dispersed intensity [periplasmic TorA (39); outer membrane BamE (R. Ieva, unpublished) (Fig. S1C-E)]. eYFP-SctV foci fluoresced more strongly than the monomeric inner membrane protein LacY-eYFP (Fig. S1F)(40), suggesting that SctV self-assembles in higher order quaternary states in the inner membrane *in vivo* (see below). Analysis of 10,000-14,000 individual cells showing distinct clusters (Fig. S2B and C), revealed that commonly ∼1-4 eYFP-SctV clusters are formed per EPECΔ*sctV*/C41 cell (Fig. S2D) and are distributed along the cellular periphery (Fig. S2E).

### SctV natively extracted from membranes is nonameric

To study the oligomeric state and assembly of SctV, we isolated EPEC membranes, treated them with various concentrations of the non-ionic detergent dodecyl-maltoside (DDM), separated the resulting complexes by blue native-polyacrylamide gel electrophoresis (BN-PAGE) and visualized them by immuno-staining with an antibody against the C-domain of SctV (Fig. 1D).

DDM concentrations ≤0.7% [w/v; 13.2 mM; ∼80-fold above the critical micellar concentration (CMC_H2O_ = 0.17 mM)] did not solubilize discrete SctV-containing complexes (Fig. 1E, lane 2). At DDM concentrations ≥1.0%, the SctV-containing species migrated as a tighter band, with an apparent mass of ∼700 kDa, consistent with the mass of a stable nonamer (Fig. 1E, lanes 3-5).

When membranes were solubilized with 0.8% DDM, higher order SctV-containing complexes with apparent sizes of ∼1 MDa immuno-stained with α-SctV-C (Fig. 1E, lane 6, bracket). These might be sub-assemblies of the injectisome [mass >3 MDa; at least 18 different proteins (8, 35)]. When 0.8% DDM extracts were treated with additional DDM to a final concentration of 1.8%, a sharp band of SctV_9_ was obtained (Fig. 1E, lane 7), suggesting that the assembled nonamer was a pre-existing component of the ∼1 MDa species. Elevated amounts of DDM likely dissociated all other partner subunits but left the SctV_9_ intact.

These data suggested that full-length SctV in EPEC membranes assembles in stable, detergent-extractable, nonameric assemblies that easily dissociate from other injectisome components depending on detergent concentration.

### Self-assembly of SctV nonamers in membranes is injectisome-independent

SctV might nonamerize on demand by enveloping existing membrane-embedded cores of SctRSTU or other injectisome components (22, 41–43). However, EPEC-derived SctV_9_ is stable at up to 1.8% DDM (Fig. 1E) and SctV_9_ forms clusters *in vivo* even in C41 that is devoid of any injectisome components (Fig. 1C). This raised the possibility that SctV might self-assemble into nonamers in membranes.

To test this, we over-expressed the *sctV* gene behind a T7 promoter in C41 cells grown in LB medium (Fig. S3A, lane 3). The membranes were isolated and the membrane- embedded SctV complexes were characterized by BN-PAGE as above. At low DDM concentrations (0.4% DDM), C41 extracts contained negligible amounts of DDM-solubilized SctV (Fig. 1F, lane 2), while at DDM concentrations above 0.6%, discrete nonameric species of SctV of ∼700 kDa were detected (Fig. 1F, lanes 3-7), similar in size to those seen in the EPEC-derived extracts (Fig. 1E, lanes 3-5). One visible difference though was that SctV_9_ extraction from C41 membranes initiated at almost half the DDM concentration needed to extract it from EPEC membranes (Fig. 1F) suggesting that in EPEC, additional components may further stabilize membrane-embedded and assembled SctV_9_. The SctV_9_ species can also be extracted with other non-ionic detergents (Fig. S3B), implying a stable, pre-existing, detergent-independent, oligomeric state. SctV_9_ in C41 membranes was destabilized and aggregated by heat treatment (Fig. S3C) or once purified from membranes, dissociated into smaller complexes when treated with an ionic detergent (Fig. S3D), but was very stable at high concentrations of different non-ionic detergents (Fig. S3E and F).

SctV_9_ may interact with additional injectisome components (16, 33). We tested directly whether the export apparatus components SctRSTU form complexes with SctV_9_. His-SctRSTU synthesized in C41, gives rise to a complex that is best extracted with 3.6% digitonin and has a mass of ∼240kDa when analyzed by BN-PAGE and immuno-stained with α-SctU C-domain antibody (Fig. 1G, lane 1). This apparent mass would be consistent with the R_5_:S_4_:T_1_:U_1_ stoichiometry anticipated from cryo-EM and MS experiments (3, 10, 16, 33, 34). When co-synthesized with SctV in the same cells, a higher order species of >1MDa is observed at the low DDM concentration of 0.8% used (Fig. 1H, lane 4). This species runs as a diffuse band suggesting poor stability and increased dynamics. Once sequentially treated with 1.8% DDM (w/v) the nonameric species of SctV of ∼700kDa appears as a prominent sharp band, indicating dissociation from other components (Fig. 1I, lane 2).

Collectively, these data suggested that SctV_9_ stably self-nonamerizes in the absence of a scaffold provided by other T3S components but weakly associates with pre-assembled SctRSTU once the latter is present in the membrane.

### Contributions of the N- and C-terminal domains of SctV to its nonamerization

SctV comprises a 34.8 kDa N-terminal domain with 8 predicted transmembrane helices connected by a short ∼3 kDa linker to a 37.1 kDa C-terminal cytoplasmic domain. To determine which SctV segment is important for nonamerization, His-tagged N-terminal and C-terminal SctV fragments (Fig. 2A, top) were expressed in C41 and their oligomerization properties were compared to those of SctV after cytoplasmic and membrane extracts were fractionated by ultra-centrifugation.

**Fig. 2.**
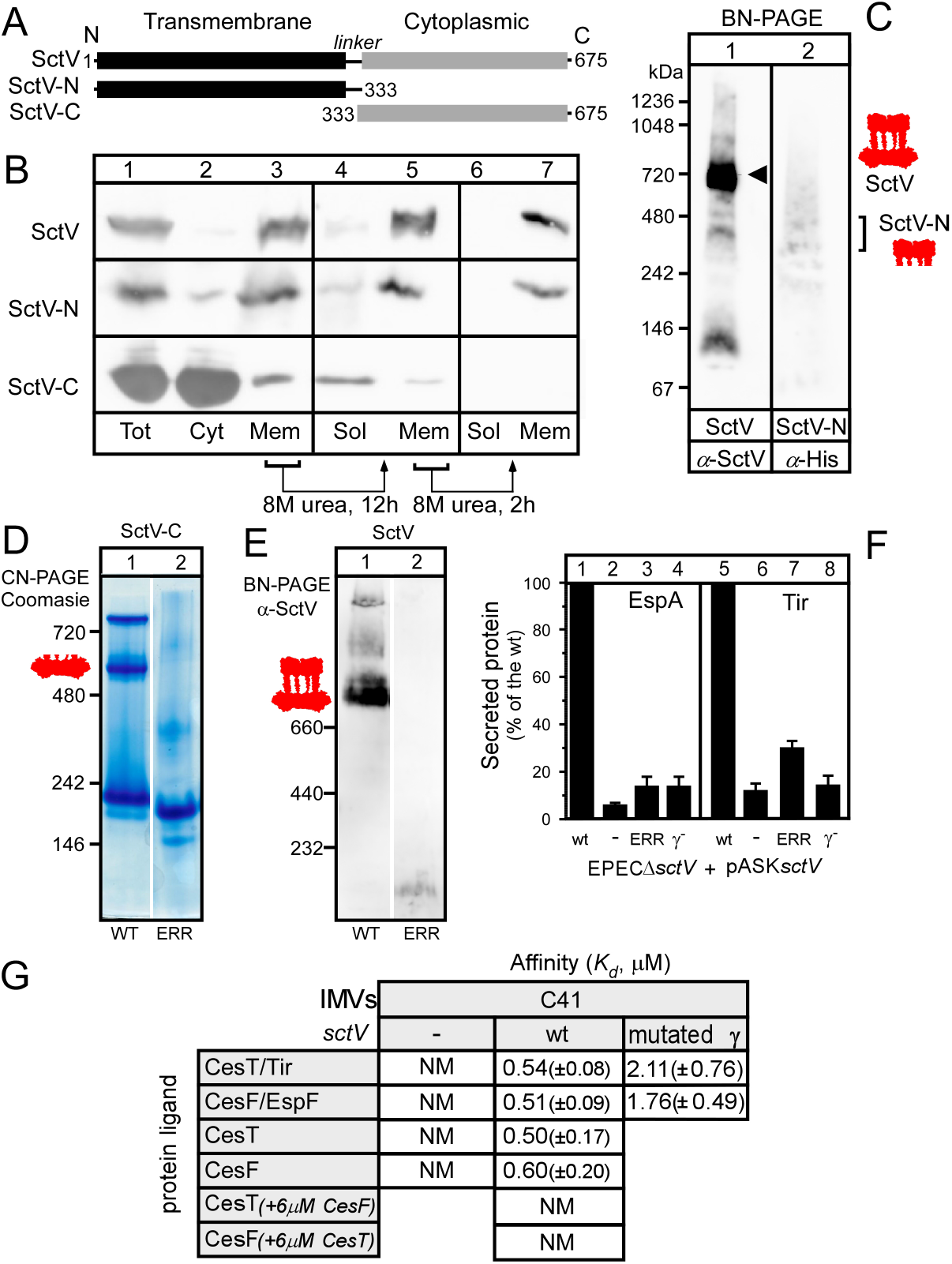
Analysis of SctV_9_ as a functional receptor for chaperone/exported protein complexes. **A.** Schematic representation of SctV transmembrane, linker and cytoplasmic domains (20) **B.** Sub-cellular localization of full length SctV and its sub-domains in C41 cells expressing plasmid-borne SctV, SctV-N or SctV-C (Bottom panel). Following fractionation, equal fraction volume was analyzed by 12% SDS-PAGE and immuno-staining. Tot: Total cell lysate; Cyt: cytoplasmic fraction; Mem: Membrane fraction; Sol: Solubilized membrane protein fraction. Representative images are shown; *n=3*. **C.** BN-PAGE analysis of His-SctV-N in C41. The membrane fraction of C41 cells expressing plasmid-borne SctV or His-SctV-N was extracted with 1.8% DDM. Samples were analyzed by BN-PAGE and immuno-staining with an α-SctV-C or α-His antibody (lanes 1 and 2, respectively). Representative image is shown; *n=3* **D.** The oligomerization states of SctV-C and its derivative. Purified SctV-C and one of its interface disrupting mutants were analyzed by CN-PAGE and stained with Coomasie. SctV- C mutant ERR: E503A-R535A-R564A; Representative images are shown; *n=3* **E.** The oligomerization states of full-length SctV and its derivative. As in C, proteins from membrane fractions derived from C41/*sctV* wt or mutated were extracted with 1.8% of DDM, analyzed by BN-PAGE, and immuno-staining with α-SctV-C. SctV mutant ERR: E503A- R535A-R564A; Representative images are shown; *n=6*. **F.** Secretion of EspA and Tir from EPEC Δ*sctV* cells carrying *sctV* wt or derivatives (as indicated). Secreted protein amounts were quantified and secretion derived from EPEC Δ*sctV*/ *sctV* wt was considered 100%. SctV mutant ERR: E503A-R535A-R564A; SctV Patch γ-: DKITFLLKKL➤GSIAVLLASS; All other values are expressed as % of this. n=3. **G.** Equilibrium dissociation constants (*K_d_*) of protein ligands for IMVs derived from C41 cells with or without SctV or with SctV mutant (as indicated). The *K_d_*s of CesT and CesF were also determined for IMVs that had been pre-incubated with excess of CesF or CesT, respectively (bottom); SctV Patch γ-: DKITFLLKKL➤GSIAVLLASS; mean values (± SEM) are shown; *n=6*.

Full-length SctV and SctV-N were mainly distributed in the membrane fraction, while most of SctV-C was detected in the cytosolic fraction (Fig. 2B, lanes 1-3). To exclude co-sedimentation with membranes due to aggregation and inclusion body formation, the membrane fraction was further washed with 8M Urea in two steps. Full-length SctV and SctV-N remained largely in the urea-insoluble membrane fraction (Fig. 2B, top and middle, lane 7), while SctV-C was extracted by urea completely (Fig. 2B, bottom, lane 7). We concluded that SctV-N can become membrane integrated in the presence or absence of the soluble C-domain. Membrane-embedded His-SctV-N extracted with detergent migrates diffusely at apparent masses of 200-500 kDa in BN-PAGE gels (Fig. 2C, lane 2, bracket), that are discrete from those of SctV_9_ (lane 1). This might imply formation of dynamic but unstable oligomers.

Next, we purified His-SctV-C by metal affinity chromatography and sucrose gradient ultra-centrifugation (Fig. S4A and B). His SctV-C had a mass of 384.3 (± 3.5) kDa and a hydrodynamic (Stokes) diameter of 15.14 (± 0.82) nm (shown by analytical gel permeation chromatography coupled to multi-angle and quasi-elastic light scattering detectors (GPC- MALS/QELS) (Fig. S4C) or 379.2 (± 2.07) kDa (by native mass spectrometry; Fig. S4D). Both in good agreement to the theoretical mass of His-SctV-C_9_ (374.9 kDa) indicating that the C-domain of SctV inherently self-nonamerizes.

To test the nonamer-promoting role of the C-domain in the context of the full-length SctV, we introduced different mutations in residues of SctV-C that lie at the protomer-protomer interface [(12, 14); see below]. In all cases, the mutations resulted in loss of the nonameric states of both the SctV-C (Fig. 2D; lane 2, one example shown; all mutants in Fig. S5A) and the SctV full length (Fig. 2E; lane 2, one example shown; all mutants in Fig. S5B). All oligomerization mutants were functionally defective in the secretion of either middle (Fig. 2F, lane 3, one example shown; all mutants in Fig. S5C) or late (lane 7 one example shown; all mutants in Fig. S5C) clients, indicating that self-nonamerization is essential for secretion and that several residues on the oligomerization surface have similar essential contributions to oligomerization.

We concluded that SctV-C has a strong tendency to self-nonamerize. This leads to nonamerization of the full-length SctV, perhaps aided by weak SctV-N oligomeric assemblies in the membrane.

### SctV_9_ is a necessary and sufficient receptor for chaperone/exported client complexes

SctV interacts with T3S chaperones with or without exported clients and/or the gatekeeper SctW (9, 29, 44). Nonamerization may be required for high affinity binding since chaperone/exported substrate complexes bind with low affinity to monomeric flagellar SctV (29). To determine whether self-assembled SctV_9_ is functional as an exported protein receptor, we probed its binding to the dimeric chaperone CesT alone or in complex with its cognate client Tir, using an *in vitro* affinity determination assay developed previously (9). IMVs were prepared from *E. coli* C41, harbouring either *sctV* on a plasmid or the empty vector and were urea-treated to strip away peripheral and non-specifically associated proteins. The external face of the IMVs corresponds to that of the cytoplasm in the cell; their lumen represents the periplasmic face. SctV_9_-containing, C41-derived IMVs were functionally competent for high affinity saturable binding of CesT or the CesT/Tir complex (Fig. 2G; Fig. S6A), with *K_d_* values equivalent to those measured in IMVs from EPEC cells in the presence of the full complement of injectisome components [Fig. 2G; (9)]. In contrast, IMVs from C41 devoid of SctV showed only a linear, non-specific binding component that did not yield a measurable *K_d_* (Fig. 2G; Fig. S6A). The receptor function of SctV was further corroborated using a second chaperone and chaperone/effector complex CesF/EspF (45) that also bound with high affinity to SctV-containing C41-derived IMVs, but not to C41- derived IMVs not containing SctV (Fig. 2G; Fig. S6B).

Chaperone/exported protein pairs are likely to share common receptor docking regions on the SctV_9_ receptor (29). Attesting to this, incubation of excess of CesF chaperone (at concentrations of 10-fold *K_d_*) prevents the binding of CesT on IMVs prepared from C41/*sctV* and *vice versa* (Fig. 2G).

Chaperone/exported middle client complexes and the SctW gatekeeper bind to six different patches on the SctV C-domain determined using arrays of immobilized peptides (9). Using the same approach CesT, CesT/Tir, CesF, and CesF/EspF were shown to bind to patches α−ε (Fig. S6C-E) that may be used in multi-valent binding (see below). IMVs prepared from C41 cells expressing *sctVγ^-^* were used as an example. SctVγ^-^ formed nonamers as did the wild type (Fig. S5D, lanes 1 and 2) but showed a 3-4-fold affinity reduction for CesT/Tir and CesF/EspF (Fig. 2G). All patches are functionally important beyond their *sensu stricto* receptor capacity for *in vivo* secretion of middle and late clients (9)(Fig. 2F, lanes 4 and 8, the γ derivative is completely defective).

We concluded that SctV_9_ embedded in C41 membranes is necessary and sufficient to act as a high affinity receptor for T3S chaperone/exported client complexes *via* its cytoplasmic domain. The important mechanistic implication of this observation is that the receptor function of SctV neither depends on the presence of any other T3S components nor on ongoing translocation.

### Purified full length His-SctV_9_ forms a ring structure

To determine the ultrastructure of full-length SctV_9_, we synthesized His-SctV in C41. His-SctV forms nonamers like SctV (Fig. S1G, lanes 1 and 2), was extracted from membranes with 1.8% Triton X-100, and separated from higher order aggregates on a sucrose gradient (Fig. 3A and B; Fig. S4E). Fractions enriched in His-SctV_9_ (determined by BN-PAGE; Fig. 3C) were collected, pooled and incubated with Ni^2+^-nitrilotriacetic acid (Ni^2+^- NTA) resin. Resin-bound His-SctV_9_ assemblies were on-bead reconstituted into peptidiscs using 30% fluorescently labelled peptides (46). To probe if the peptidisc particles were indeed reconstituted in soluble states in the absence of detergent, we analysed them by clear native PAGE (CN-PAGE), which separates molecules on the basis of charge, mass and shape. Silver-stained CN-PAGE gels of His-SctV_9_-PR revealed a band of ∼700-800 kDa (Fig. 3D) that was also fluorescent, indicating the presence of the bound solubilizing peptidisc peptides (Fig. 3E). GPC-MALS/QELS revealed that His-SctV_9_-PR has a mass of 762.8 ± 6.9 kDa (Fig. 3F, red) consistent with a nonameric species bound to ∼17 solubilizing peptides of 4.5 kDa and a hydrodynamic diameter of 22.0 ± 0.98 nm (Fig. 3F, blue) close to the anticipated dimensions derived from cryo-ET (16).

**Fig. 3.**
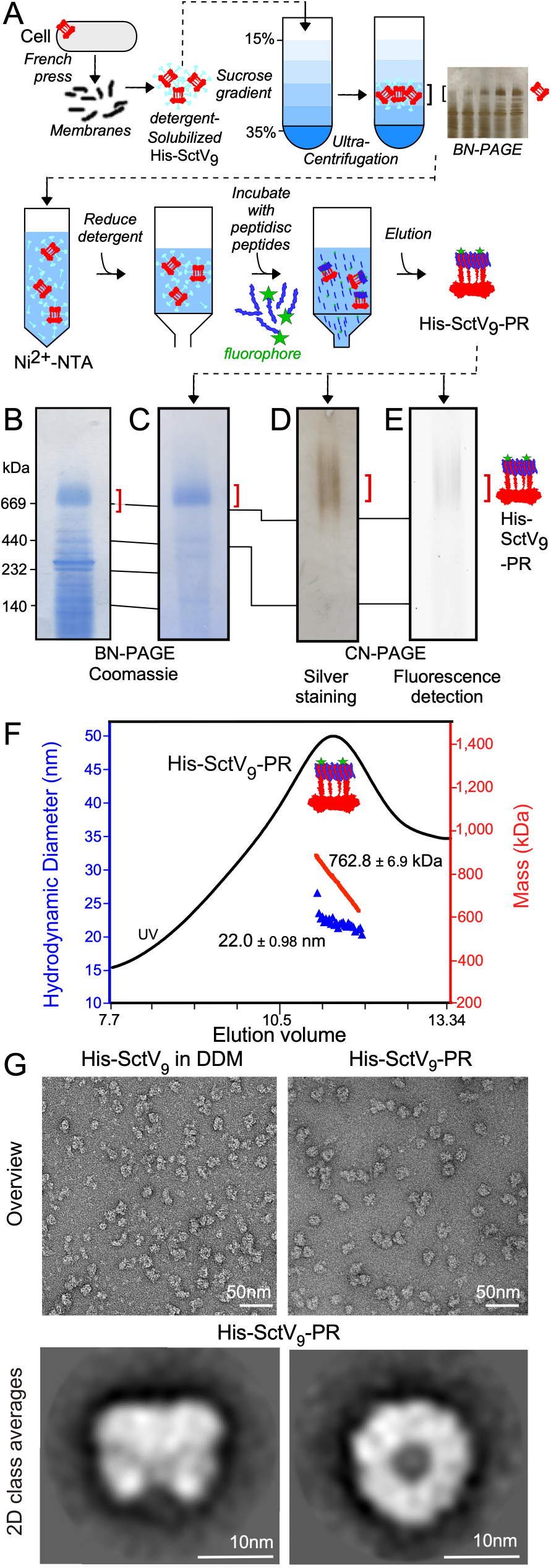
Full-length SctV purification, peptidiscs reconstitution, and characterization. **A.** Pipeline of removal of SctV higher order aggregates and on-bead reconstitution into peptidiscs (His-SctV_9_-PR) and purification. Crude membrane fraction of C41/*his-sctV_9_* cells, following French press disruption and ultra-centrifugation, was treated with 1.8% v/v Triton X-100 and loaded onto the 15%-35% sucrose gradient to separate His-SctV_9_ from soluble higher order aggregates. **B.** Sucrose gradient fractions containing His-SctV_9_ without higher order aggregates were analyzed by BN-PAGE and stained with Coomassie blue (representative image shown). These fractions were used for further peptidisc reconstitution. Representative image is shown; *n=3* **C-E.** Validation of His-SctV_9_-PR reconstitution. His-SctV_9_-PR was analyzed by BN-PAGE followed by Coomasie blue staining (**C)** and CN-PAGE followed by silver staining (**D**) and fluorescence peptidisc detection **(E).** Representative images are shown; *n=3*. **F.** GPC-MALS/QELS analysis of His-SctV_9_-PR using Superose^TM^ 6 (GE Healthcare). UV (in black), mass traces (in red) and hydrodynamic diameter (in blue) of are shown. *n=3*. **G.** Negative staining EM analysis of His-SctV_9_-PR. Representative “overview” (upper panel; scale bar: 50 nm) and “side” and “bottom views” of 2D class averages from His-SctV_9_-PR are shown (lower panel; scale bar: 10 nm).

The His-SctV_9_-PR was analysed by negative staining EM (Fig. 3G, overview). Class averaged side views of individual His-SctV_9_-PR particles (Fig. 3G, bottom left) reveal that they comprise two distinct domains likely representing the membrane-embedded N-domain and cytoplasmic C-domain parts of SctV separated by a narrow constriction (9, 12, 16). Class averaged bottom views reveal a multimeric ring with an inner opening (Fig. 3G, bottom right).

### High resolution structure of the SctV C-terminal domain nonameric ring

Repeated attempts to obtain a high-resolution structure of full length SctV failed to yield sufficient detail in the membrane part of the ring. We therefore focused our attention on the cytoplasmic domain. His-tagged SctV-C was purified and its structure determined using cryo-EM. Both single- and double-ringed species were observed in an approximately 4:1 ratio and were sorted out by using a 3D classification in Relion 3.1 (Fig. S7A; Table S1). The reconstructed maps of the single-ringed particles with imposed C9 symmetry and double-ringed particles with imposed D9 symmetry were refined to 3.6 Å and 3.4 Å resolution, separately (Fig. S7A). The map of the single-rings (map1) is well aligned with parts of the double ringed map (map2) (Fig. S7B). Therefore, the double-ringed particles were subtracted into two single-ringed particles and combined with the single-ringed particle population. Thus, a final 3.1 Å map (map3) using 105,670 particles from 3739 micrographs was achieved (Fig. S7C-D). Local resolution regions of the EM map are consistent with predicted disordered regions (Fig. S6E). The model of the regions of 345-415, 462-583, and 626-670 of SctV-C was obtained *de novo* from map3. While, the initial model of _Sf_SctV-C_9_ (PDB: 4a5p, 2x49) (12, 20) was used to guide the model building process for _EPEC_SctV-C (Fig. 4A).

**Fig. 4.**
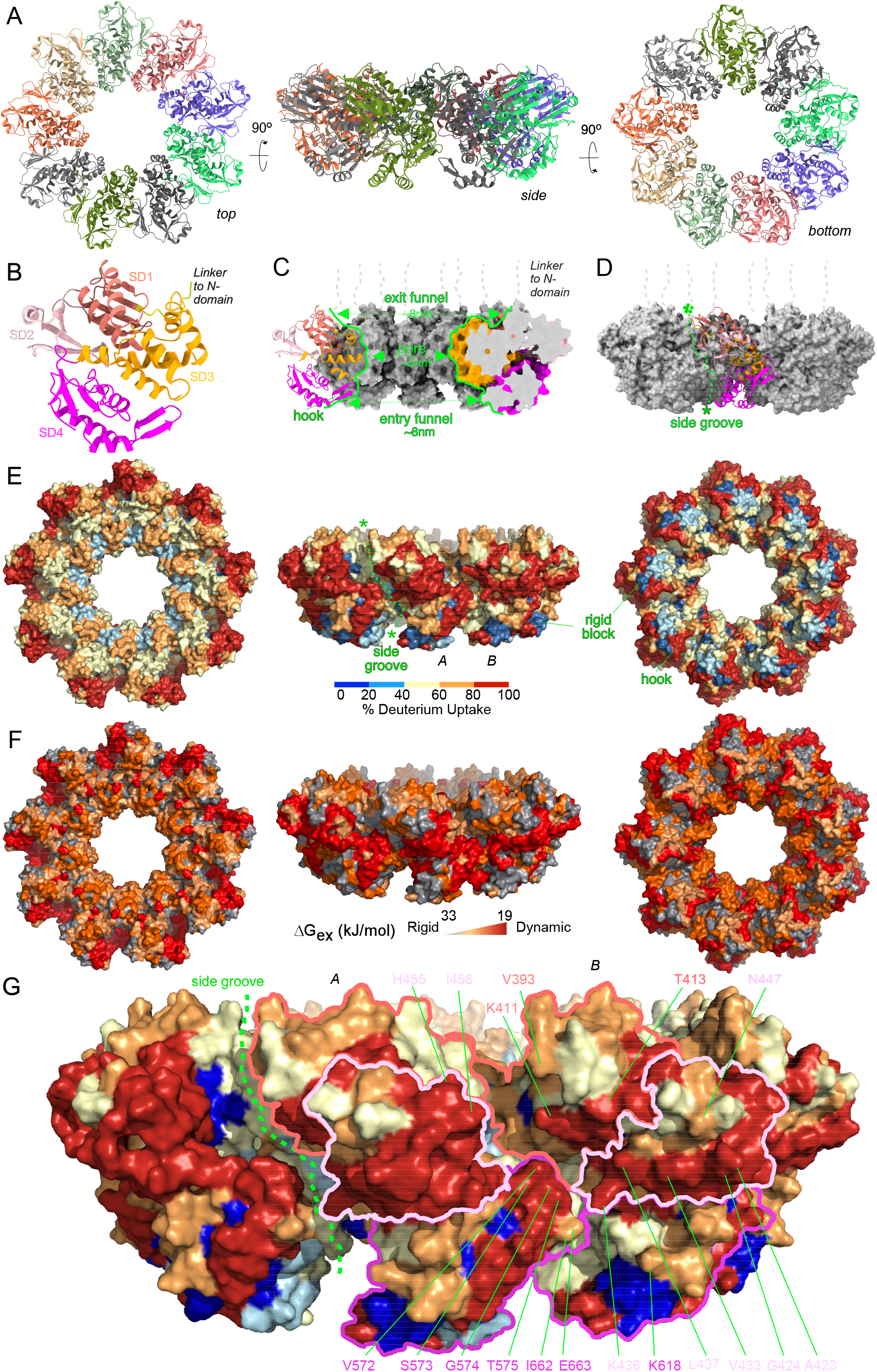
Structure and structural dynamics of SctV-C_9_ using Cryo-EM and HDX-MS. **A.** Overviews of the nonameric SctV-C model. Different views of SctV-C_9_ model were shown with each protomer being differentially colored. **B.** Ribbon diagram indicating the sub-domains of SctV-C (SD; SD1: 353-415 and 463-483; SD2:416-463; SD3: 488-570; SD4: 570-676). The protomer is shown in side view orientated as the orange protomer in 4A, side view. **C.** Central cross-section showing the two-funnel-like transport channel of SctV-C. Two cross sectional protomers were shown in ribbon and surface mode, and coloured as in panel B. The entry/exit funnels, the central pore, and the hook-shape region of SctV-C are indicated with green arrows. **D.** Side groove formed by two adjacent protomers of SctV-C. The groove of one protomer (shown in ribbon and colored as in panel B) is indicated with a green line. Asterisks indicate the entry and exit site. **E.** Local intrinsic dynamics of SctV-C_9_ assessed by HDX-MS. SctV-C_9_ indicated views in surface representation, colored according to the %D uptake. Differences (D) in %D at 5 min are visualized on the structure of SctV-C_9_ using a color gradient that correlates with the % of D uptake relative to the fully deuterated control (as indicated). Side groove is indicated with a green line. Asterisks indicate the entry and exit site. Blue hues: rigid regions; Red hues: disordered regions; grey: unidentified. Biological replicates *n=2,* Technical replicates *n=3*. **F.** ΔG_ex_ values (in kJ/mol) were calculated for each residue of SctV-C by PyHDX (51) from time-course D-exchange HDX-MS experiments and visualized on the nonameric SctV-C protein structure. Residues are colored on a linear scale from grey (33 kJ/mol, rigid) to red (19 kJ/mol, dynamic). Highly rigid residues (ΔG_ex_ >33 kJ/mol, transparent grey). **G.** As in E, SctV-C_9_ side view in surface representation, colored according to the %D uptake is shown, highlighting flexible regions on one protomer. Charged residues of those regions that are important for interactors binding are indicated, colored according to the subdomains that they belong to.

The SctV-C_9_ ring contains a charged pore lined with positively charged residues as are the two outer rims, close to the entry and exit mouths of the pore, at the cytoplasmic-facing bottom and the membrane-facing top of the structure, respectively (Fig. S8A). The outer periphery of the ring is characterized by separated upper and bottom negatively charged bands. Significant conservation is seen in the residues that build the nonamer and line the inner ring pore (Fig. S8B). In contrast, outwardly facing peripheral surfaces of the ring are not conserved.

The SctV-C protomer is connected to the transmembrane N-domain by a linker (Fig. 2A; 4B; S6E) and is built of four sub-domains (SD) connected by hinges. SD1 is discontinuous (aa353-415 and 463-483); SD2 (416–463) is inserted in SD1; SD3 (488-570 and SD4 (571–676) [Fig. 4B; Fig. S6E and S8D (47)]. SD3 is the only domain that lines the narrowest constriction of the pore and nonamerizes the protomer and is the most conserved domain, as are some regions of SD1 and SD4 (Fig. S8B).

The nonamer assembles using an interprotomeric interface that is not extensively hydrophobic and includes multiple intermolecular electrostatic bridges that connect one sub- domain in one protomer A to a sub-domain in the adjacent protomer B, primarily through conserved residues in SD3-SD3 and SD3-SD1 contacts. These connect linker_A_-SD3_B_ (S347-R532), SD1_A_-SD3_B_ (E483/R564) and SD3_A_-SD3_B_ (Q488-E503/Y501 and E489-R535) interfaces (Fig. S8C-G). Moreover, the SD1-SD3 hinge, is positioned between two protomers and may contribute to ring formation and conformational changes (47). All of these residues, except E503, were shown by mutagenesis to be crucial for nonamerization of both SctV-C and full length SctV (Fig. 2D and E; S5A and B; S8F and G) and for secretion (Fig. 2F; S5C). The oligomerization interface, provided primarily by SD3 residues (Fig. S8D), is the most conserved part of the protein (Fig. S8B). In contrast, the regions that are away from the ring pore formed by the conserved SD3, including the outwardly facing SD1 and 2 and parts of SD4 are the least conserved.

The ring pore is built exclusively of SD3 contributed from the 9 subunits (Fig. 4C), with the highly conserved K506_SD3_, R510_SD3_, and K549_SD3_ residues lining it (Fig. S8E) and has a diameter of ∼5nm. The cytoplasmic entry to the pore is built exclusively of SD4 that lines the pore entry with its 2β-tip (residues 594-604; β11/12) that forms a hook-like protrusion (Fig. 4C). SD4 is recessed outwards relative to SD3, thus providing a wider, funnel-like entry mouth to the pore (Fig. 4C, entry funnel) with a diameter of ∼8nm between hooks. On the other end, the exit mouth of the pore facing the membrane, widens out away over SD3, is built exclusively by SD1 and together with the top of SD3 form a wide exit mouth “ledge” (Fig. 4C, exit funnel). SD2 is unrelated to the pore and faces outwards. When a protomer is viewed head on in a side view of the ring, SD3 and SD4 have a near vertical placement, like barrel staves, while SD1/2 tilt leftward towards the adjacent subunit by ∼30° from the vertical axis (Fig. 4D). This positioning creates large elongated grooves between the staves (Fig. 4D, dashed line) including the conserved residues R532_SD3_, E407_SD1_ and R564_SD3_ (Fig. S8B, middle) and also on either side of the ring “castle embrasures”, between adjacent SD4 domains on the entry funnel at the bottom and between SD1 domains at the exit funnel at the top(asterisks).

### Local structural dynamics of SctV-C_9_ determined by HDX-MS

Given that the same nonameric SctV ring changes its affinities for clients on the basis of associated protein regulators (9), we hypothesized that the intrinsic local dynamics of SctV may underlie its function. To test this, we used a hydrogen deuterium exchange mass spectrometry pipeline (HDX-MS; Fig. S9) which non-invasively monitors loss or gain of backbone H-bonds, commonly participating in secondary structure. The labeling reaction is at low micromolar concentrations and at physiological buffers (48–50). SctV-C was diluted to ∼5 µM into D_2_O buffer for various time-points. Samples were acid-quenched, protease-digested and D-uptake was determined. 233 peptides with good signal/noise ratio yielded ∼99.7% primary sequence coverage (Table S2). The D-uptake for a tandem array of peptides that cover all of the sequence was expressed as a percentage of each one’s fully deuterated control (taken as 100%) (Fig. 4E). Dynamics data are subsequently used to derive Gibbs free energy of exchange (ΔG_ex_, kJ mol^-1^) per residue of a protein’s sequence using our in-house software PyHDX (Fig. 4F)(51, 52). ΔG_ex_ is inversely correlated to the degree of dynamics in the protein backbone, therefore lower ΔG_ex_ values represent higher backbone flexibility.

SctV-C displays significant dynamics including regions yielding lower resolution in the cryo-EM structure and predicted to contain disordered sequence (Fig. S6E; S7D). The highest level of dynamics is observed in the periphery of the ring (Fig. 4E and F, left and middle) and the hook at the entry funnel (right). In contrast, the pore and the exit funnel are rather rigid (Fig. 4E and F, left) as is a region next to the hook (Fig. 4E, middle and right). The observed elevated dynamics of the ring align along the ridges that flank the side groove, in which one elevated dynamics region of each protomer apposes the elevated dynamics region of the adjacent protomer (Fig. 4G).

### Intrinsic local dynamics of the SctV-C protomer

To gain insight in the regions of elevated dynamics and rigidity in the four sub- domains of the SctV protomer we mapped the HDX-MS data (Fig. 4E) onto a protomer structure derived from cryo-EM in ribbon representation (Fig. 5A). Highly dynamic residues are in red hues, while the less dynamic ones are in blue hues. SD2 displays the most elevated and SD3 the more reduced dynamics in the structure. The only regions of SD3 that display elevated dynamics are its connecting linker to SD4 (Fig. 5A.I, V572-T575) and a short connection between α5 and α6 (G513-I517) that faces the inner ring surface (Fig. 5A.II). On the other hand, the hinge that connects SD3 to SD1 shows reduced dynamics (III). In addition to its intrinsic flexibility, SD2 is connected to SD1, into which it is rooted, using highly flexible loops connecting to the highly flexible ends of two secondary structure elements (β4 and α3; IV) and this would greatly facilitate rigid body motions of SD2 around this two-element hinge. The highly flexible β4 of SD1 participates in a 4-stranded sheet, displaying additional flexible regions (V), that face the side groove of the ring (Fig. 4G). SD4 is rather rigid with three main internal regions of high dynamics: the hook (β10-β11; VI), the loop_N604-R609_ (VII) and the extreme C-terminal region (I662-A676; VIII). These overall features were largely retained in the SctV monomer, obtained by analysing the monomeric SctV(R535A) derivative. Some regions of the monomer displayed more enhanced (e.g. V477-L504 in SD3 that face the pore) and two more reduced (N345-L357 and D550-L554) dynamics (Fig. 5B). These differences were visualized in a differential uptake map were purple hues indicated enhanced rigidity and green hues enhanced dynamics (Fig. 5C).

**Fig. 5.**
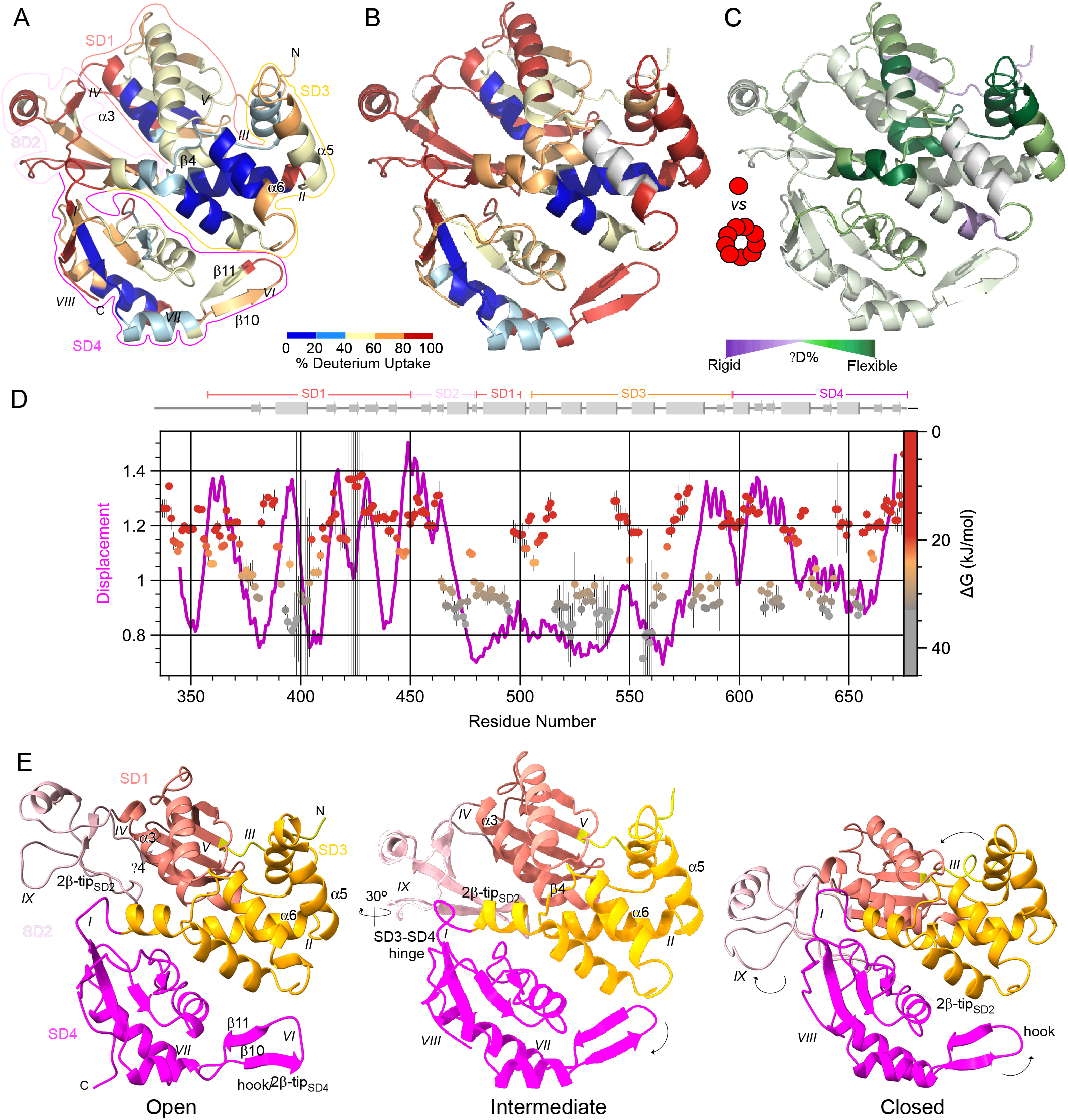
Local and domain dynamics in the SctV-C protomer. **A.** Conformational dynamics of the SctV-C protomer assessed by HDX-MS and colored in a ribbon representation (as in Fig. 4E). Tracing lines indicate the respective subdomains, colored as in Fig. 4B; Latin numerals indicate important dynamic motions; Secondary elements are represented for clarification. **B.** Conformational dynamics of the SctV-C R535A mutant assessed by HDX-MS and colored in a ribbon representation (as in Fig. 4E). %D uptake is shown. **C.** Differences (D) in %D uptake on mutating residue R535A (which forms monomer) at 5 min are visualized on the structure of SctV-C9 using a color gradient that correlates with the % of D uptake relative to the fully deuterated control (as indicated). Highly flexible regions are displayed in green and rigid regions are shown in purple on mutating R535A. **D.** ΔG_ex_ values were calculated for each residue of SctV-C using PyHDX software (as in Fig. 4F). ΔG_ex_ values are shown per residue of SctV-C-domain (residues 334-675) and colored on a linear scale from grey (33 kJmol^-1^; rigid), to orange (21 kJmol^-1^, flexible) to red (19 kJmol^-1^; dynamic). Normal modes are calculated using the WebNM@ web serve. Displacements per normal mode are then summed with equal weights for the first 6 non-trivial normal modes. Normal mode flexibility is derived from normal mode eigenvalues as described previously. **E.** Comparison of SctV-C sub-domains in the three indicated states derived from analysis of the 2 PDB structures integrated into a movie (Supplementary methods; Movie S1). The _EPEC_SctV-C structure determined here is that of the “intermediate” state. Important elements and arrows showing domain movements are indicated.

This pattern of intrinsic dynamics of SctV-C yields several regions of the protein with residues that have very low ΔG thresholds between open and closed states, explaining their flexibility (Fig. 5D, coloured circles). This pattern of flexibility was also corroborated by an orthogonal approach, normal mode analysis, in which low frequency vibrational states for the atoms of a protein can be derived using coarse grain models derived from PDB structures and calculating their possible displacement (Fig. 5D, magenta line) (53). In the context of the nonamer-embedded protomer, flexible regions are located exclusively in SD1,2 and 4 and are practically absent from SD3.

### Sub-domain motions in SctV-C

Local dynamics in critical parts of a structure, as detected in Fig. 5A, B and D for SctV-C, are commonly accompanied by and permit domain motions (14, 27, 52). All available structures of SctV-C have been solved in slightly different domain arrangements revealing a repertoire of domain conformational states that can be acquired by the four sub- domains of the C-domain. These can be seen to broadly ranging from an “open” to a “closed” state (12, 14, 18, 27, 29), primarily determined by the motion of the hyper-flexible SD2. The SctV-C structure resolved here approaches the closed state but its SD2 is closer to its SD1 compared to other structures and its SD2 is more detached from SD4 than in the “closed” models. To visualize the inherent dynamics of the C-domain sub-domains in the context of the protomer and the nonamer, we modeled the potential transition motions from an open to a closed state of SctV-C homologues into a movie (Fig. 5A; Movie S1).

The main rigid body motions modelled are those of the SD2 which makes use of its hinge connecting it to SD1 (linkers between β4 and β5 and β8 and α3; Fig. 5A.IV; E, left; S6E) to extend outwards from the ring periphery to yield an “open” state and then retracts to an “intermediate” position, resting its 2β-tip_SD2_ (residues 433-444; β6/7) against SD1 and a flexible loop (residues 423-429; IX) against the C-terminus of SD4. This retraction of SD2 is coincident to a rigid body rotation of SD4 around its hinge with SD3 by ∼30° (Fig. 5E, middle). This brings the protruding hook/2β-tip_SD4_ (β10/11; VI) to move towards SD3 and the inside of the pore. In a second step, SD1 undergoes a rigid body motion along its hinge with SD3 (Fig. 5E, right, III) and moves towards SD4. This displaces the 2β-tip_SD2_ which now moves towards the C-terminus of SD4, while the 423-429 loop of SD2 (IX) moves away from SD4. Concomitantly, the hook reverses its previous motion (Fig. 5E, right; Movie S1).

During the “pinching” motion of SD2 against SD4 in the ring periphery to yield the closed state, the inner ring surface and the diameter of the pore remain largely unchanged, except for individual charged and bulky residues of the inner ring side of SD3 (aa K500, N502, E503, K506, F543, K549; Fig. S8E).

Collectively, this modelling suggested possible rigid body domain transitions derived from the high-resolution structural snapshots of end states. Our HDX-MS data would now rationally explain these motions as resulting by the local dynamics of StcV-C and involving the hinges and the hyper-flexible SD2 (Fig. 5A; Movie S2). Rather than distinct stable “states”, the intrinsic dynamics analysis raises the more possibility of a conformational ensemble of states.

### SctV-C sub-domain dynamics are related to association of interactors

Two main groups of SctV interactors have been analysed to date: chaperones with or without clients (24, 29, 47, 54) and the inner stalk subunit SctO of the ATPase complex (27). Binding of chaperone/client complexes occurs mainly in the periphery of the ring extending to the exit funnel. Binding sites determined by NMR and peptide arrays are located on either side of the side grooves of the ring (Fig. 6A; S6E; Movie S3). A prominent chaperone binding site (patches α and β; S6E) is the 4-stranded beta sheet of SD1 that forms a continuous surface with SD2 residues, with residues like (V393_SD1_, K411_SD1_, T413_SD1_, K436_SD2_, L437_SD2_ in protomer B and apposed to H455_SD2_, I456_SD2_ in protomer A; Fig. 6B, top) overlooking the side groove (Fig. 4F). The non-conserved and highly dynamic SD2 is a major site of chaperone binding on a single protomer but can also contribute binding interfaces for multivalent inter protomeric binding. Sub-domain motions affect the degree of exposure of the binding sites (Movie S3). Most patches and NMR binding sites also coincide wholly or partly with islands of high or very high flexibility, particularly on SD2 and SD1 (Fig. S6E). Mutagenesis of surface-exposed residues in several of these regions revealed that they all contribute to chaperone binding affinity and are all essential for secretion [patch γ as an example; Fig. 2F, lanes 4 and 8 and 2G;(9)].

**Fig. 6.**
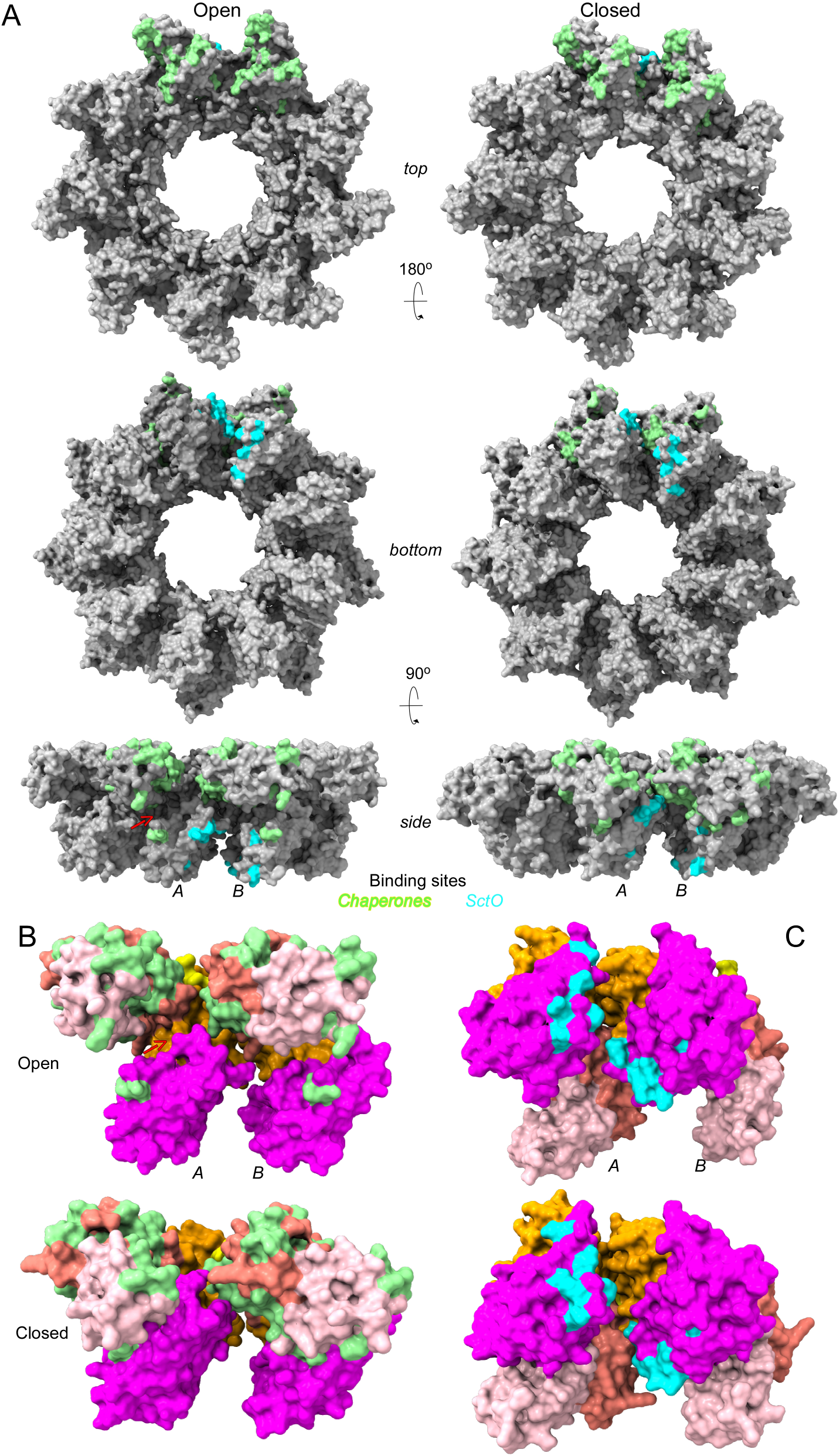
Chaperone and ATPase inner stalk binding sites on the SctV-C protomer. **A.** Chaperone and SctO binding domains in the indicated views of the SctV-C_9_ surface structure at the open (left) and closed (right) state. Chaperone binding sites determined from peptide arrays (Fig. S6E, α,β and γ) and NMR analysis (9, 29) are coloured light green in two adjacent protomers (dark grey) of the nonamer model. SctO binding sites determined from cryoEM analysis (27) are colored cyan. For detailed residues and mutations see Fig. S6E and Table S3. Red arrow: crevice between SD2-SD4 revealed by SD2 opening. **B.** Chaperone binding sites (light green) are shown in the open (top) and closed (bottom) state SctV-C of two adjacent protomers (coloured as in Fig. 4B) in a surface representation. Red arrow: crevice between SD2-SD4 revealed by SD2 opening. **C.** SctO binding sites (cyan) from (27) are shown on open (top) and closed (bottom) state SctV-C of two adjacent protomers (coloured as in Fig.4B) in a surface representation.

SctO binds exclusively to SD4, almost vertically against the entry funnel of the ring [Fig. 6A;(27)] approaching it from the cytoplasm and inserting in sockets formed by the vertical walls of two adjacent SD4 domains and a flat roof provided by the bottom side of SD3 (Fig. 6 C; Movie S4). SctO binds on one side directly the SD4 hook of protomer A, and on the other SD4 residues of protomer B.

## DISCUSSION

How injectisomes function remains elusive. With the development of *in vitro* reconstitution systems (9, 55), guidance from cryo-EM-derived structures and structural dynamics introduced here for the first time, this issue can now be systematically addressed, and assembly processes can be dissected away from functional ones (i.e. secretory client docking and translocation).

The assembly of the SctRSTU translocase and its SctV export gate are key to the initiation of the downstream export events (22, 42). A major mechanistic challenge for the assembly of all these subunits is that although they enter the membrane using the universal membrane-protein integration machineries (*e.g.* Sec/SRP, YidC), SctV needs to subsequently multimerize and be “plugged” by SctRSTU. While oligomerization in the presence of other injectisome components *in vivo* has been implied for *_Ye_*SctV-eGFP (43), our experiments demonstrate that SctV self-assembles into functional nonamers, driven largely by its C-domain (Fig. 2C-D; S5A; S8), *in vivo* and *in vitro* even in cells that have no injectisome components. Therefore, nonamerization is an inherent SctV property. In the absence of SctV, the rest of the injectisome still assembles and SctRSTU acquires its periplasmically-exposed location inside the SctD/J rings (16, 22). Thus, SctV, perhaps pre-assembled, may arrive late to half-pre-assembled injectisomes into which it can integrate to meet SctRST (16). In EPEC, this may be simply a kinetically controlled stochastic event, through tight regulation of transcription and/or translation, given that SctV is synthesized in cells much later than SctRSTU (56).

SctV-C/FlhA-C form nonameric rings in solution via electrostatic interactions (14, 34, 57). Here with an improved map at 3.1 Å resolution we confirmed that multiple interfaces (Fig. S8C-G) are indeed involved in cytoplasmic ring assembly. Mutational analysis revealed that these contacts are all essential in forming the nonameric ring both in the isolated C-domain (Fig. 2D; S5A) and in the full-length protein (Fig. 2E; S5B), as is the interprotomeric SD1-SD3 hinge involved in conformational changes (47)(Fig. S8C-G). Formation of the C-domain ring also leads to nonamerization and function of the full-length SctV, with a minor possible contribution from the transmembrane N-domain. The latter domain cannot be resolved well by cryo-EM in isolated SctV_9_ (17, 21) presumably due to the absence of other T3SS components and/or lipids.

A significant body of data had implied that SctV possesses significant dynamics: the loss of nonamerization by single residue C-domain mutations (Fig. S5A-B), the reduced resolution in regions of the cryo-EM map, alternate domain states in the different SctV structures (12, 14, 47), the significant effects of patch mutations in both the affinity and the secretory function of SctV (Fig. 2F, G; S5C), the isolation of point mutants in SctV that mimic the effects of bound interactors (58), the ability of the ring to switch affinities (9) and be influenced by pH (17) and Ca^2+^ ions (14). The local HDX-MS analysis performed here provided for the first-time residue level structural understanding of these dynamics (Fig. 4E; 5A). It revealed the coexistence of a very rigid pore-building/nonamerizing sub-domain attached to two moderately dynamic domains that build the entry and the exit funnels of the pore. Moreover, it revealed a hyper-dynamic, non-conserved, outward facing sub-domain that controls chaperone access to the ring periphery. Flexible and inflexible hinges between the sub-domains and flexible elements like the entry funnel hook and the inner pore exposed dynamic region of SD3 (Fig. 5A) suggest possibilities for motions and functional roles. In common with multiple other examples (59–61), the dynamics described here are intrinsic, and reflect an inherent property of the monomeric polypeptide (Fig. 5B). As such, their exploitation by binding ligands does not require high energy input.

Our data corroborated and extended the major role of SctV as a chaperone/exported client receptor (9, 29), via its C-domain. T3SS chaperones are structurally distinct and flexible and atomistic detail of their binding in the context of the SctV nonamer is missing. Nevertheless, localization of binding interfaces on the SctV-C_9_, hints to important functional regions made available between adjacent protomers, in the context of the nonamer (Fig. 6A and B, light green; Movie S3). Several residues in these regions are important for chaperone/gatekeeper binding in structures solved by NMR (29) and in genetic analyses (23, 24, 62–64)(Fig. S6E; green; Table S3). Using different T3SS-chaperones we demonstrated that binding sites on SctV-C are universal and shared (Fig. 2G). The absence of conservation, charged surfaces with excessive dynamics would allow the hyper-flexible SD2 to co-evolve to specifically bind the multiple different T3SS chaperones of a particular organism. The rather flat nature of this receptor landscape and its dynamics make it structurally feasible to accommodate very large chaperone/client complexes, while dimeric chaperones and/or clients may additionally secure docking on multiple regions of adjacent protomers. NMR structures suggested that chaperone complexes insert their binding elements inside both the side groove of the ring but also in the crevice between SD2-SD4 uncovered when SD2 moves outward to the open state [(29); Fig. 6B, red arrow]. SD2 interactions with chaperones may also control their release or accessibility since chaperone binding sites are hidden when SD2 retracts (Movie S3, green) (14, 27).

The identified chaperone binding regions on SctV-C have important mechanistic implications as they demonstrate that while targeting and subsequent secretion are structurally and biochemically separable, these processes must also be somehow coupled since when the receptor site γ is mutated alone, it leads to a modest reduction in affinity but completely abrogates secretion [Fig. 2F-G; (9)].

Multiple limited area binding sites on receptors, dynamic non-folded polypeptides and excessive dynamics are common themes in chaperone-non-folded client recognition and delivery systems (52, 65, 66). Multi-valent binding secures synergistic low nM affinities, from binding sites that individually possess only μM binding strength and the non-folded client does not face a severe entropic penalty when bound as it retains most of its sequence unstructured. A powerful mechanistic reason for these evolutionary choices is their facile loss of the nM binding for the chaperone and/or client and the release of the latter by minor conformational modulation of the receptor. This “disruptive” role is anticipated here to be played by the SctO inner stalk of the ATPase since it penetrates between two adjacent SctV- C subunits at SD4 and binds the hook [Fig. 6C; (27)] and will undoubtedly affect the intrinsic dynamics of both subunits and the wider ring. Moreover, SctO binding might stabilize the SD2 open state [MovieS4; (27)]. As the machinery has nine binding sites for SctO and chaperone/clients but only one SctO subunit attached to one hexameric ATPase, a rotary mechanism would disrupt consecutively bound multiple chaperone/client complexes. Because the receptor is intrinsically dynamic, the ATP expenditure for client release may be minor.

Future analysis will be necessary to dissect the structural basis of these events and define the path that the exported clients follow upon their dissociation from their chaperones. As SD2 retraction coincides with SD4 hook rotation towards the central pore, this motion of and those of highly conserved pore-lining residues K506_SD3_, R510_SD3_ and K549_SD3_, might also guide and translocate clients into the pore (Fig. S8 D and E; Movie S1). Both the hook and the conserved pore-lining residues affect substrate switching (17). The development here of tools like the peptidisc-reconstituted SctV_9_ pave the way for functional reconstitution of the inner membrane translocase to decipher translocase mechanism and dynamics.

## Materials and Methods

For the complete list of strains, plasmids, mutants, primers, buffers, antibodies, see the supplementary material.

### Cell growth, induction of gene expression, *in vivo* secretion

EPEC strains were grown in the optimized M9 medium (67) (37°C; 6 h). Plasmid gene expression was induced from the tetracycline promoter (OD_600_=0.3; AHT; 2.5 ng/mL; 3 h or as indicated). Cells were harvested (5,000 x g; 20 min; 4°C; Sigma 3-16KL; rotor 11180); the spent medium was TCA-precipitated (20% w/v) and resuspended with 50 mM Tris-HCl (pH 8.0) in volumes adjusted according to OD_600_. An equal number of cells or supernatant volumes derived from an equal number of bacterial cells, were analyzed by SDS-PAGE, and immuno- or Coomasie Blue- stained.

### Live-cell imaging

Cells were imaged on inverted fluorescence microscope (Olympus IX-83) with a 1.49 NA oil-immersion objective (Olympus UAPON 100x) and ET442/514/561 Laser triple band set filtercube (69904, Chroma). The excitation light was provided by a 514 nm laser (Sapphire LP, Coherent) and the fluorescence emission was collected on an EM-CCD Camera (Hamamatsu C9100-13). Camera frames were collected using CellSens software (Olympus) for multi-position acquisition. Brightfield images were flattened by subtracting dark offset and dividing by an empty brightfield image. Fluorescence image were corrected for illumination beam profile and dark count. The brightfield images are segmented to identify cell positions by convolutional neural networks with the U-Net architecture (68–70). Next, individual cells were analyzed with the software ColiCoords (71). Binary objects from segmented images were filtered based on morphological features (size, ellipse axis) to select only single planktonic *E. coli* cells. Next, the obtained cell objects were further filtered in consecutive steps 1) Shape of the binary image; 2) radius of the cell measured from the brightfield image (C41; 280 nm ⩽ r ⩽ 437 nm, EPEC; 467 nm ⩽ r ⩽ 568 nm); 3) shape of the brightfield image radial distribution; 4) fluorescence intensity, which resulted in removal of out-of-focus cells and overexpressing cells. To localize fluorescent foci, first the cytosolic background was obtained by filtering the images with a spatial median filter (72) (kernel size 5 pixels). Foci were identified in the background-subtracted images by a local maximum filter (73). To remove aggregates and inclusion bodies from the identified peaks, the Zernike moments (74) for each peak were determined [inline], and by thresholding only Gaussian-shaped peaks were selected.

### Membrane solubilization and BN-PAGE analysis

Cell pellets were resuspended in Buffer A supplemented with 1 mM MgCl_2_, 2.5 mM PMSF, and 50 mg/mL DNase and lysed by using French press (1,000 psi; 6 passes). The unbroken cells were removed by centrifugation (3,000 x g; 4°C; 5 min; Sigma 3-16KL; rotor 11180). The membrane fraction was pelleted down by high-speed centrifugation (100,000 x g; 4°C; 30 min; 45Ti rotor; Optima XPN-80, Beckman Coulter). The membrane pellet was resuspended in solubilization buffer, homogenized with a Dounce homogenizer to a final protein concentration of ∼40 mg/mL. Membrane proteins were extracted by incubating with non-ionic detergent as indicated (4°C; 1 h). Following ultra-centrifugation (100,000 x g; 4°C; 15 min; rotor TL-100; OptimaMax-XP, Beckman-Coulter), solubilized membrane protein samples (20μl extract with BN-PAGE loading buffer, 10% glycerol) were loaded on 3-12% gradient Bis-Tris NativePAGE^TM^ precast protein gels (Invitrogen) and subjected to BN- PAGE electrophoresis as described previously (75) with little modification. BN-PAGE electrophoresis was performed with anode buffer and cathode buffer 1 using XCell^TM^ *SureLock*^TM^ Mini-Cell (approximately 2.5 hours, 4°C, 100V) until the blue running front has moved about one-third of gel. Then cathode buffer 1 was replaced with cathode buffer 2 for electrophoresis (15 hours, 4°C, 70V).

### Reconstitution of His-SctV_9_ in peptidiscs

The peptidisc reconstitution of His-SctV was conducted ‘on-bead’ as described previously (46) with some modifications. Crude membrane suspensions were derived from C41 cells overexpressing His-*sctV* by French press (6 passes; 1,000 psi; pre-cooled with ice). A total of 10 mL suspension (200 mg/mL) was mixed with 200 μl 100 x protease inhibitor cocktail solution (Sigma Aldrich, Cat. S8830), solubilized in Buffer F (supplemented with 1.8% Triton X-100) and incubated (1 h; 4°C). Insoluble aggregates were removed by centrifugation (20,000 x g, 30 min, 4°C; Sigma 1-16K). Solubilized membrane proteins (1 mL; total of 400 mg) were loaded on a 30 mL 15-35% w/v linear sucrose gradient prepared as previously described (76). Sucrose solutions were made in Buffer G and 25 x 89 mm 38.5 mL open-up polyallomer tubes were used (Beckman Coutler). Ultra-centrifugation was performed (160,000 x g; 16 hours; 4°C; SW 32 Ti rotor; Optima XPN-80, Beckman Coulter) and 1mL fractions were collected and analyzed by BN-PAGE. Fractions containing SctV_9_ without higher order aggregates of SctV were selected and incubated with 1.5 mL of Ni^2+^- NTA resin (Qiagen) pre-equilibrated in Buffer F (6 h; 4°C). The Ni^2+^-NTA beads were transferred to a gravity column, washed with 100 column volumes (CV) of Buffer F supplemented with 0.9% Triton X-100 and 10 CV of Buffer H supplemented with 0.02% DDM, sequentially. Post-washing, 1 CV of Assembly Buffer containing 0.8 mg/mL NSPr mix (1 fluorescent: 2 non-fluorescent NSPr; Peptidisc, an amphipathic bi-helical peptide; PEPTIDISC BIOTECH) in 20 mM Tris-HCl pH 8.0) was added to the beads and incubated (2 h; 4°C). Following washing with buffer H (10 CV), His- SctV_9_-PR was eluted in 0.5 mL fractions (5 mL of Buffer H + 300 mM imidazole). The elution fractions were loaded onto a Superose^TM^6 10/300 GL column previously equilibrated with buffer L using an ÄKTA Pure system (GE Healthcare). The purified His-SctV_9_-PR Peptidisc complex was subsequently analyzed by negative-stain electron microscopy.

### Purification of His-SctV-C_9_

BL21(DE3) cells overexpressing *his-sctV-C* were resuspended in buffer A and lysed by French press 1,000 psi; 5-6 passes; pre-cooled with ice). Insoluble material was removed by centrifugation (100,000 x g; 30 min; 4°C; 45 Ti rotor; Optima XPN-80; Beckman Coulter). The cell lysate supernatant was passed through a Ni^+2^-NTA resin column (Qiagen), and protein was purified following the manufacturer’s instructions. Eluted His-SctV-C was loaded on a 13 mL 15-35% linear sucrose gradient and centrifuged (as described above). Fractions were loaded on 3-12% gradient Bis-Tris NativePAGE^TM^ precast protein gels (Invitrogen) and subjected to CN-PAGE electrophoresis. CN-PAGE electrophoresis was performed using XCell^TM^ *SureLock*^TM^ Mini-Cell (approximately 15 hours, 4°C, 15 mA) and gels stained with Coomasie Blue. Fractions containing mainly His-SctV-C_9_ were further purified on a Superdex200 10/300 GL column previously equilibrated with buffer M using an ÄKTA Pure system (GE Healthcare). His-SctV-C_9_ complexes were subsequently analyzed by cryo-EM.

### Negative Staining and EM

Sample preparations with the concentration of 20-100 ng/μl were applied to glow-discharged, carbon-coated copper grids, and stained with a 2% (wt/vol) solution of uranyl acetate, and examined under a FEI Talos L120C (120 KV) microscope equipped with a 4k × 4k Ceta^TM^ 16M camera. The micrographs were acquired at 92,000X magnification with 2 seconds exposure. Particle picking and 2D classification were performed using Relion 3.1(77)

### Single-Particle cryo-EM and Data processing

SctV-C was vitrified on Quantifoil 1.2/1.3. Briefly, 4 μl of the sample was applied onto glow discharged grids and allowed to disperse for 0.5-2min. The grids were blotted for the 4-7s set at 100% humidity and plunge-frozen in liquid propane/ethane mixture cooled with liquid nitrogen to about minus 180 -190 °C by using a Vitrobot Mark V. Vitrified specimen were imaged on a FEI Titan Krios operating at 300 kV and equipped with a field emission gun (XFEG) and a Gatan Bioquantum energy filter. Movies consisting of 25 frames were automatically recorded using FEI EPU software and the K2 Summit camera resulting in 0.55 Å per physical pixel. For individual frames, an electron dose of 1.65 e^-^/Å^2^ was used, corresponding to a cumulative electron dose of 41.25 e^-^/Å^2^ equally distributed over a 5-sec movie. Movies were recorded at 0-4 µm defocus. Samples for diameter measurements were recorded with LEGINON13 on a FEI Polara (300 kV) equipped with a field emission gun (FEG) and a Gatan CCD Camera (UHS 4000). Single-particle reconstructions were performed using Relion 3.1 (77). 3739 Movies were motion-corrected, dose-weighted, and binned by 2. The CTF estimation of the resulting micrographs was determined using CTFFIND4 (78). Particles were picked from the motion-corrected micrographs using crYOLO (79) trained with a sub-set of manually picked particles. Particles were extracted into 256 x 256 boxes and subsequently binned by 2 for several rounds of 2D classification. A cleaned dataset was obtained by re-extraction and aligned to a rotationally averaged structure. 3D classification with C1 symmetry was performed to sort out the single-ringed particles and double-ringed particles. Focused refinements with applying C9 for single- ringed particles and D9 symmetry for double-ringed particles were performed. After converged refinements, per-particle CTF and Bayesian polishing were used to generate new data sets for another round of focused refinements. Overall gold-standard resolution: Fourier shell correlation (FSC=0.143) and local resolution were calculated with Relion3.1. The double-ringed particles were subtracted into two single-ringed particles and combined with single-ringed particles for the refinements for generating a better single-ringed map.

### Model building, refinement and validation

Ab initio model building was performed with Coot (v0.9-beta)(80). Interactive refinement against the cryo-EM map density was performed with ISOLDE (v.1.1.0)(81), a molecular dynamics-guided structure refinement tool within in ChimeraX (v.1.1)(82). The resulting coordinate file was further refined with Phenix.real_space_refine (v.1.18-6831)(83) using reference model restraints, strict rotamer matching and disabled grid search. Model validation was carried out using MolProbity server (84) and EMRinger (85) within the Phenix software package. More details can be found in Table S1.

### SctV sub-domain motions analysis

_EPEC_SctV-C was compared to its homologs with existing PDB files using ChimeraX 1.0 (https://www.rbvi.ucsf.edu/chimerax/). The conformational states were then classified into different classes according to the distance between SD2 and SD4. Two models were generated after structural alignment using chimeraX 1.0 that had apparently the most extreme distances in the conformational states of SD2 and SD4. To build the open and close state of the _EPEC_SctV-C structural model we used SWISS-MODEL (https://swissmodel.expasy.org/) and PDB: 6wa6 *Chlamydia pneumoniae* and PDB: 2x49 *Salmonella typhimurium* were chosen as the template for open and close state respectively. Then the open and close state of SctV-C, and the intermediate state of SctV-C obtained from cryo-EM here were morphed using ChimeraX to see how the sub-domains might move. The movie was recorded in ChimeraX and further processed using iMOVIE (https://www.apple.com/imovie/).

### HDX-MS experimental procedure, data collection and analysis

#### D exchange reaction and Quenching

The purified SctV-C_9_ protein was diluted to 5 μM for mass spectrometry experiments using H_2_O Buffer (25 mM Tris pH 8; 25 mM KCl). For HDX-MS experiments, 4 μl of SctV-C_9_ in H_2_O buffer were mixed with 46 μl of the D_2_O buffer (25 mM Tris pD 7.6; 25 mM KCl) at 30°C and incubated at various time intervals (10s, 30 sec, 1min, 5 min, 10 min, 30 min, 100 min) before being added to 50 μl of quench solution (8M Urea; 0.1% DDM; 5 mM TCEP pH 2.5). A 100-µL Hamilton syringe was used for sample injection (100 μl). The mobile-phase flow paths were held at 0.5 °C using a three-valve unit (Trio Valve, Leap Technologies) constructed with a custom thermal chamber such that on-line protein-digestion, peptide desalting and reversed-phase HPLC separation are performed prior to infusion into the ESI ion source of the mass spectrometer. Loading of sample, digestion, and desalting (3 min) was driven with an isocratic HPLC pump (IPro-500, IRIS Technologies, Lawrence, KS) at a flow rate of 100 µL min-1 through the 50-µL sample loop, the immobilized nepenthesin column (Affipro 2.1 mm ID x 20 mm length, Cat.No.: AP-PC-004), across a VanGuard C18 Pre-column, (130 Å, 1.7 mm, 2.1 x 5 mm, Waters), and out to waste. After isolation of the enzyme column from the flow path, gradient elution from the trap across a C18 analytical column (130 A°, 1.7 mm, 1 x 100 mm, Waters) was carried out with a capillary-scale HPLC pump (Agilent 1100, Palo Alto, CA). The flow rate was held constant at 40 µL min-1 while the composition of the mobile phase was increased from 0% ACN containing 0.23% (FA) to 40% ACN containing 0.23% FA over 12 min gradient. Following gradient elution, mobile- phase composition was increased over a 1-min period to 90% ACN, 0.1% FA and held at that composition for 6 min prior to reduction over 1 min to 100% H_2_O, 0.1% FA. All HPLC connections were made with 1/16 in. × 0.05 in. PEEK tubing.

#### Peptide identification and HDX data analysis

To map the peptides of SctV-C_9_, identification MS runs were performed, where 5 μM of protein in H_2_O buffer was used. The sample was quenched as described above and analysed in the MS^E^ acquisition mode in a SYNAPT G2 ESI-Q-TOF mass spectrometer over the m/z range 100-2,000 Da with the collision energy was ramped from 15 to 35 V.

HDX-MS experiments were performed on SYNAPT G2 ESI-Q-TOF mass spectrometer using a capillary voltage 3.0 kV, sampling cone voltage 20 V, extraction cone voltage 3.6 V, source temperature 80°C, desolvation gas flow 500 L/h at 150°C. The primary sequence of SctV-C_9_ as a search template was used to identify SctV-C_9_ peptides identification using ProteinLynx Global Server (PLGS v3.0.1, Waters, UK). HDX experiments were then analyzed in Dynamx 3.0 (Waters, Milford MA) software. All the other parameters were as previously described (52). Full deuteration controls were obtained by incubating SctV-C_9_ in D_2_O buffer overnight at 30°C. D-uptake (%) was calculated using the full deuteration control D-uptake values. The data has not been corrected for back exchange and is represented either as absolute D values or as a percent of the full deuteration control (86).

### Gibbs free energy and normal mode analysis

Gibbs free energy of local unfolding (ΔG_ex_, kJ mol^-1^) was determined assuming the Linderstrøm-Lang model of H/D exchange (87), using the PyHDX software (51). Normal modes of SctV were calculated using the WebNM@ web server (53), using the SctV cryo- EM structure as input (PDB XXX). Per-residue displacement was averaged across all protein chains and the first 6 non-trivial normal modes were summed with equal weights to obtain the final NMA displacement.

### Miscellaneous

*In vivo* protein secretion, IMVs preparation, peptide arrays, SEC-MALS- QELS analysis and determination of equilibrium dissociation constants (*K_d_*) were as described (9).The details of peptide array and *Kd* determination can be also found in supplementary methods. Ni^2+^-NTA purification was according to the manufacturer’s instructions (Qiagen). Purification of plasmids, PCR, digestion, and DNA fragments were done using Wizard® DNA Clean-Up System from Promega. DNA fragment ligation was done by using T4 DNA ligase as per the manufacturer’s protocol (Promega). Sequencing of the genetic constructs was performed by Macrogen Europe. Oligos were synthesized by Eurogentec.

## ACCESSION NUMBERS

PDB: 7OSL

EMD-13054

## Authors’ contributions

B.Y. performed *in vivo* and *in vitro* assays, molecular cloning, protein purification, GPC- MALS-QELS, reconstituted SctV in peptidiscs and performed negative staining, and cryo-EM data processing; A.G.P. performed molecular cloning, membrane binding assays, peptide array analysis, GPC-MALS, *in vivo* and *in vitro* assays; R.P. and S.K. performed HDX-MS experiments and analysis; Y.L. and J.S. performed live-cell imaging; J.S. performed and supervised free energy calculations of the HDX-MS data and developed analysis software; J.W. performed negative staining and cryo-EM data collection and processing, and analyzed data; D.F. built the atomic model; H.S.D. set up peptidisc reconstitution; B.S. performed native MS; M.S.L contributed in molecular cloning, antibody preparation, gene knockouts. S.K. trained, designed and supervised protein purification, detergent extraction and reconstitution, biochemical and biophysical experiments. T.C.M. supervised electron microscopy and structure determination experiments and analysis, C.G.K and T.C. contributed to data analysis, A.E., A.G.P., and B.Y. wrote the first draft of the paper with feedback from all the authors. A.E., T.C.M. and S.K. conceived and supervised the project.

## Acknowledgements

We thank K. C. Tsolis for preliminary SctV C-domain characterization and S.Krishnamurthy for help with analysis of HDX-MS data; C. Robinson (TorA-GFP), and R. Ieva (pDR*bamE- gfp)* for constructs. Work in our labs was supported by grants (to AE and TCM): T3RecS [#G002516N; Fonds Wetenschappelijk Onderzoek (FWO); https://www.fwo.be/ and “I 2408- B22” from the Austrian Science Fund (FWF); https://www.fwf.ac.at/en/]; (to AE): DIP-BiD (#AKUL/15/40-G0H2116N; Hercules/FWO); RUN (#RUN/16/001 KU Leuven); the FWO/F.R.S.-FNRS “Excellence of Science - EOS” programme grant #30550343); PROFOUND (Protein folding/un-folding and dynamics; W002421N; WoG/FWO) and (to AE and SK); C1 (FOscil; KU Leuven). This project was supported by funds through the Behörde für Wissenschaft, Forschung und Gleichstellung of the city of Hamburg available to TCM. B.Y. and Y.L. were Chinese Scholarship Council doctoral fellows. JHS is a PDM/KU Leuven fellow. The funders had no role in study design, data collection and analysis, decision to publish, or preparation of the manuscript. Image data processing has been performed at the German Electron Synchrotron Centre (DESY) using the High-Performance Computing Cluster.

## Data Availability

All relevant data are within the manuscript and its Supporting Information files and are fully available without restriction. Coordinate and associated volume and metadata have been deposited in the PDB and EMDB respectively. SctV-C EMD-xxx / PDB xxx

