## Supplementary Information for "Structural dynamics of the functional nonameric Type III translocase export gate"

#### Abbreviations

| | |
|---|---|
| AHT: | Anhydrotetracycline |
| BN-PAGE: | Blue native PAGE |
| CN-PAGE: | Clear native PAGE |
| DDM： | *n*-Dodecyl-*β*-D-maltopyranoside |
| EPEC: | Enteropathogenic *E. coli* |
| GPC-MALS/QELS: | Gel permeation chromatography coupled to multi-angle light scattering and quasi-elastic light scattering detectors |
| IMVs: | Inverted Membrane Vesicles |
| IPTG: | Isopropyl β-D-1-thiogalactopyranoside |
| T3S: | Type 3 Secretion |
| T3SS: | Type 3 Secretion System |
| TCA: | Trichloroaetic Acid |
| Triton: | Triton X-100 |

### Table of contents

#### Supplementary Figures:

**Fig. S1:** SctV derivatives can restore Type III secretion in EPECΔ*escV* (related to Figures 1 and 3)

**Fig. S2:** eYFP-SctV production, foci formation, and distribution in C41 or EPEC Δ*sctV* (related to Figure 1).

**Fig. S3**: Stability of the SctV nonameric complex (related to Figures 1 and 3).

**Fig. S4:** Purification of $SctV_9$ and SctV-$C_9$ (related to Figures 2, 3 and 4)

**Fig. S5:** Analysis of SctV oligomerization mutants (related to Figure 2).

**Fig. S6:** Binding of chaperone/exported client complexes to full-length, membrane-embedded SctV and to an immobilized array of SctV peptides (related to Figures 2, 5 and 6).

**Fig. S7:** Cryo-EM pipeline and structure analysis of SctV-C (related to Figure 4)

**Fig. S8:** Intraprotomeric domain organization/motions in the SctV-C protomer and close-up views of the interproteomeric interaction in the SctV-C nonamer (related to Figures 2, 4 and 6).

**Fig. S9** HDX-MS pipeline and HDX-MS analysis of $_{EPEC}$SctV-C (related to Figures 4 and 5).

#### Supplementary Tables:

**Table S1:** Cryo-EM data and model refinement

**Table S2:** HDX-MS data

**Table S3:** SctV residues important for chaperone binding (related to Figure 6 and S6)

#### Supplementary movies

**Movie S1:** Sub-domain motions of SctV-C

**Movie S2:** Highly dynamic regions of SctV-C determined by HDX-MS analysis.

**Movie S3:** Chaperone binding sites mapped on one protomer of $SctV_9$

**Movie S4:** Binding sites of the ATPase inner stalk protein SctO mapped on two adjacent protomers of $SctV_9$

#### Supplementary materials:

**Table S4:** Buffers

**Table S5:** Antisera

**Table S6:** Bacterial strains

**Table S7:** Vectors and genetic constructs

**Table S8:** List of primers used for gene cloning

#### Supplementary methods:

- Preparation of cells for live-cell fluorescence microscopy
- Western Blot analysis
- Sub-cellular protein localization of SctV
- Peptide array analysis
- Determination of equilibrium dissociation constants ($K_d$)
- SctV subdomain motions analysis

#### References

#### Supplementary Figures

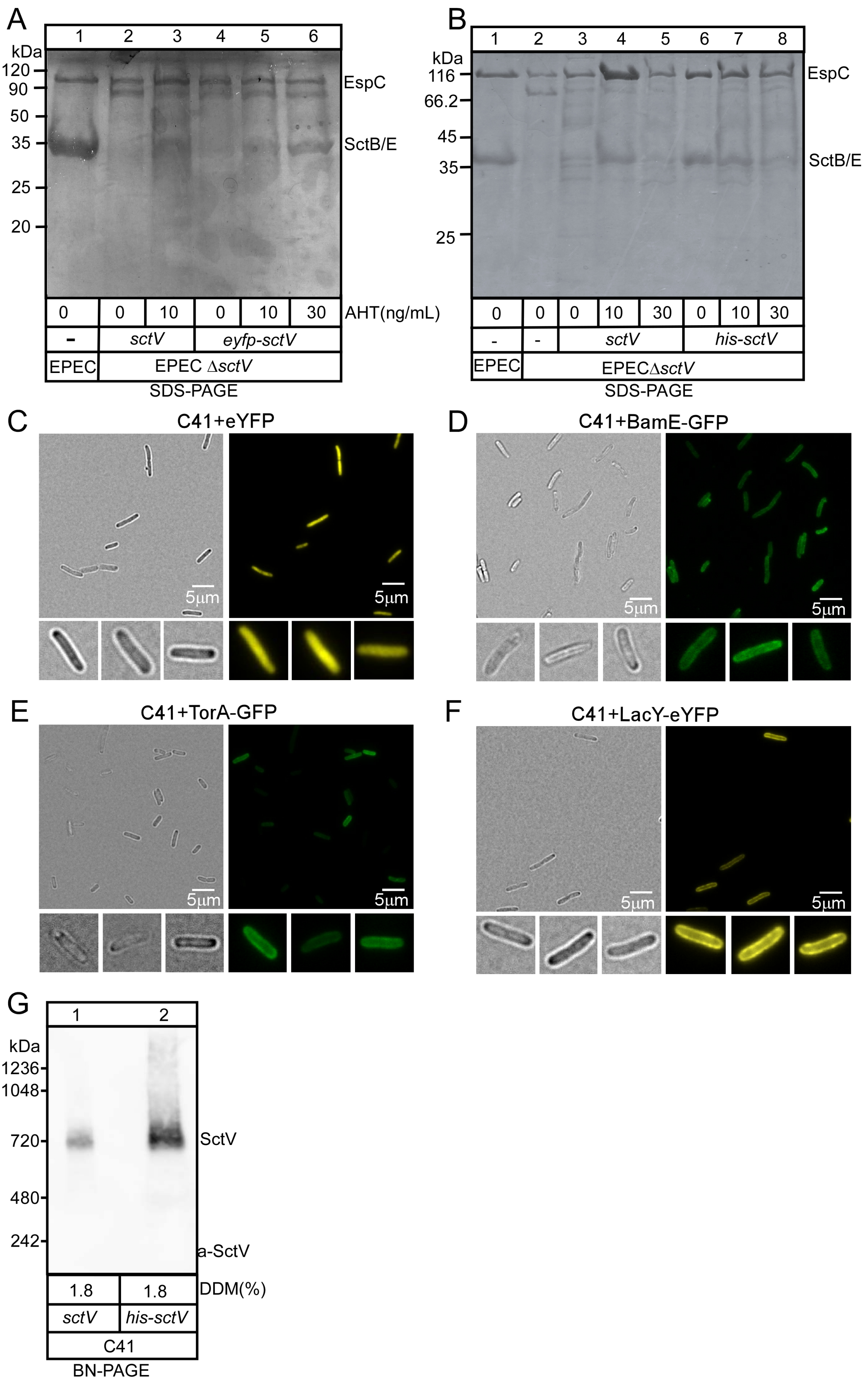

**Fig. S1: SctV derivatives can restore Type III secretion in EPECΔ*escV* (related to Figures 1 and 3).**

**A and B.** *In vivo* secretion of EPECΔ*sctV* carrying a plasmid with either *sctV*, or *eyfp-sctV* **(A)** or *his-sctV* **(B).** Gene expression was induced with different AHT concentrations as indicated. Secreted proteins from spent growth medium were TCA-precipitated. Equal volume of spent medium corresponding to equal number of cells were analyzed by SDS-PAGE, followed by coomassie blue staining. The EspC and the translocators SctB/E were indicated. Representative images are shown; *n=3.*

**C-F.** The distribution of control protein markers in C41 cells. The fluorescence micrographs of C41 cells expressing eYFP (**D**), TorA-GFP (**E**), BamE-GFP (**F**), and LacY-eYFP (**G**) separately. Scale bar: 5μm. Representative images from YFP or GFP (as indicated) and brightfield channel are shown; n=3 biological replicates

**G.** SctV and His- SctV can form nonameric species. The membrane fractions derived from C41 cells carrying pACYCDuet-*sctV* (lane 1) or *his-sctV* (lane 2) were treated with 1.8% DDM, analyzed by BN-PAGE and immuno-staining with $\alpha$-SctV-C. Representative images are shown; *n=3.*

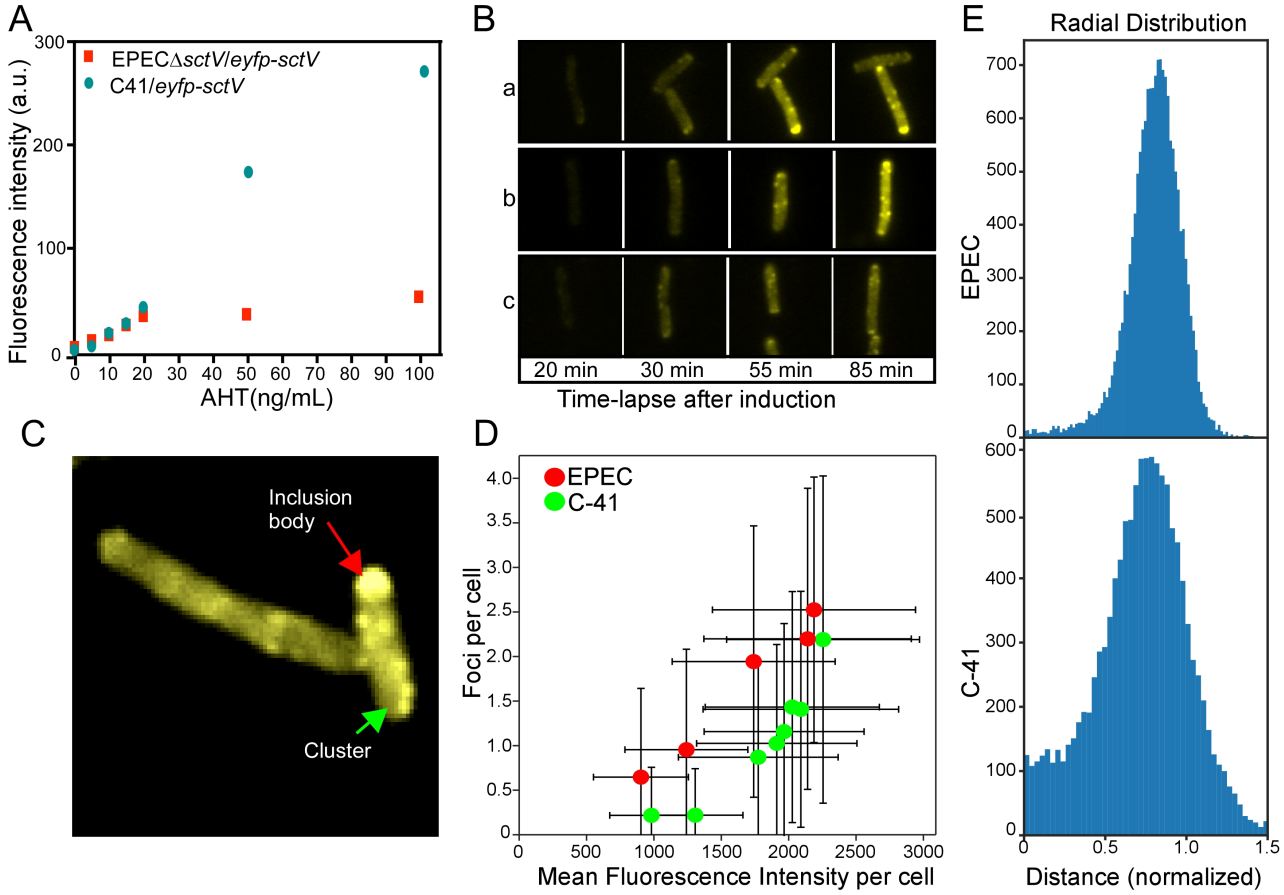


**Fig. S2: eYFP-SctV production, foci formation, and distribution in C41 or EPECΔ*sctV* (related to Figure 1).**

**A.** Comparison of the *eyfp-sctV* expression levels in C41 cells (blue) and EPECΔ*sctV* strain (red). Bacterial cells were grown in LB and induced (37°C; $OD_{600}$: 0.3) with different AHT concentrations for 3 hours. The fluorescence intensity of total cells was measured (Infinite 200 PRO microplate reader; TECAN) and used as an indicator of *eyfp-sctV* expression level. 20 ng/mL AHT was used for further experiments so to achieve similar protein production in both strains. *n=3* biological replicates

**B.** Foci formation in C41 cells expressing *eyfp-sctV*. Time lapse images were acquired every 5 min for 90 min upon induction of *eyfp-sctV* expression with 20 ngr/ml AHT. Three slices (a, b, c) are shown at different time points as indicated. *n=3* biological replicates

**C**. Inclusion body formation in C41 cells expressing *eyfp-sctV*. Inclusion body (red arrow) was distinctly larger and polarly located as compared to normal *eyfp-sctV* clusters (green arrow) and was excluded from statistical analysis.

**D**. Quantification of fluorescent foci in EPEC and C41 cells. Number of detected foci per cell for EPEC and C41 cells expressing *eyfp-sctV* were plotted as a function of the mean fluorescence intensity per cell (arbitrary units). Each data-point represents an individual microscopy experiment, error bars are standard deviations. *n=5-7* biological replicates

**E**. Foci distribution across cells. Radial distributions of fluorescent foci in EPEC (upper panel, *n=10,770*) and C41 (bottom, *n=14,209*) cells. Radial distances were measured from the center of the cell and normalized with respect to the cell's radius. Cell radius was measured from the brightfield image.

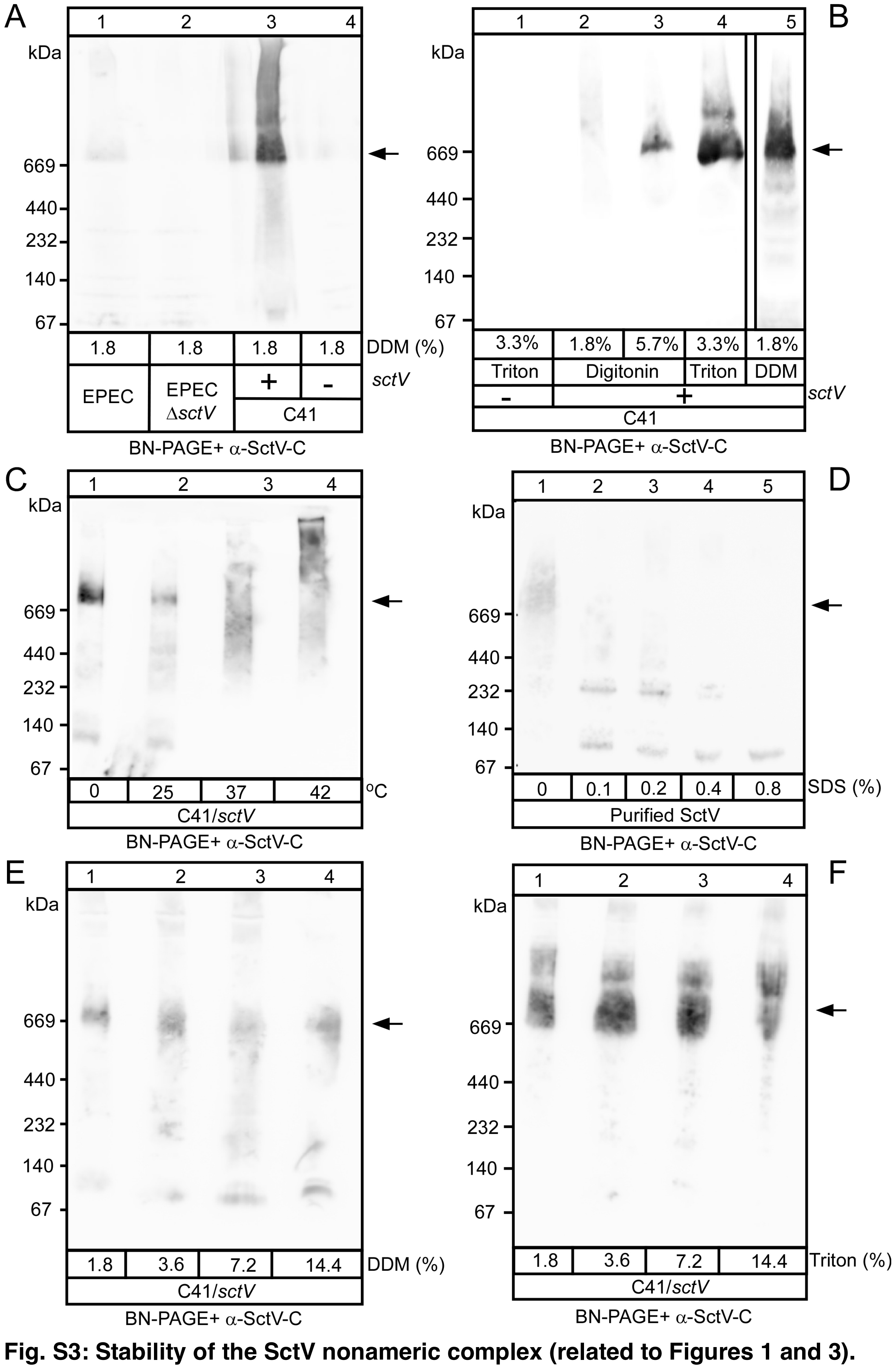


**Fig. S3: Stability of the SctV nonameric complex (related to Figures 1 and 3).**
**A**. Comparison of the *sctV* expression in EPEC (lane 1) and C41/pACYCDuet-1-*sctV* (lane 3). Membrane fractions were extracted with 1.8% DDM and samples were analyzed by BN-

PAGE and $\alpha$-SctV-C immuno-staining. 30 µl of EPEC derived membrane extracts and 5 µl from C41 (40mg/ml) were loaded on gel. Signal intensities were quantified using Image J software. >50-fold more SctV synthesis is observed in C41/pACYCDuet-1-*sctV* compared to that in EPEC; Representative images are shown; *n=3. Arrow:* $SctV_9$.

**B**. The $SctV_9$ species can be extracted with different non-ionic detergents. Membrane extracts (20 µl loading; 40mg/mL membrane protein) from C41 carrying the pACYCDuet-1-*sctV* were treated with the indicated non-ionic detergents and analyzed by BN-PAGE and $\alpha$-SctV-C immuno-staining. Representative images are shown; *n=3*.

**C.** High temperature causes $SctV_9$ aggregation. Membrane fractions were extracted (as in A) with 1.8% Triton X-100 and incubated at different temperatures (as indicated; 30 min). Samples were analyzed by BN-PAGE (20 µl loading; 40mg/mL) and $\alpha$-SctV-C immuno-staining. Representative images are shown; *n=3.*

**D.** Ionic detergent (SDS) disassembles $SctV_9$. Purified His-SctV (2 µg; 200 ng/µl) was treated with different amounts of SDS (as indicated; 30 min; 4°C) and then analyzed by BN-PAGE and $\alpha$-SctV-C immuno-staining. Representative images are shown; *n=3*.

**E** and **F.** $SctV_9$ is resistant to high concentration of non-ionic detergents. The membrane fractions of C41/*sctV* were extracted with different high concentrations of DDM (**E**) or Triton X-100 (**F**) and then analyzed by BN-PAGE and $\alpha$-SctV-C immuno-staining. Representative images are shown; *n=3.*

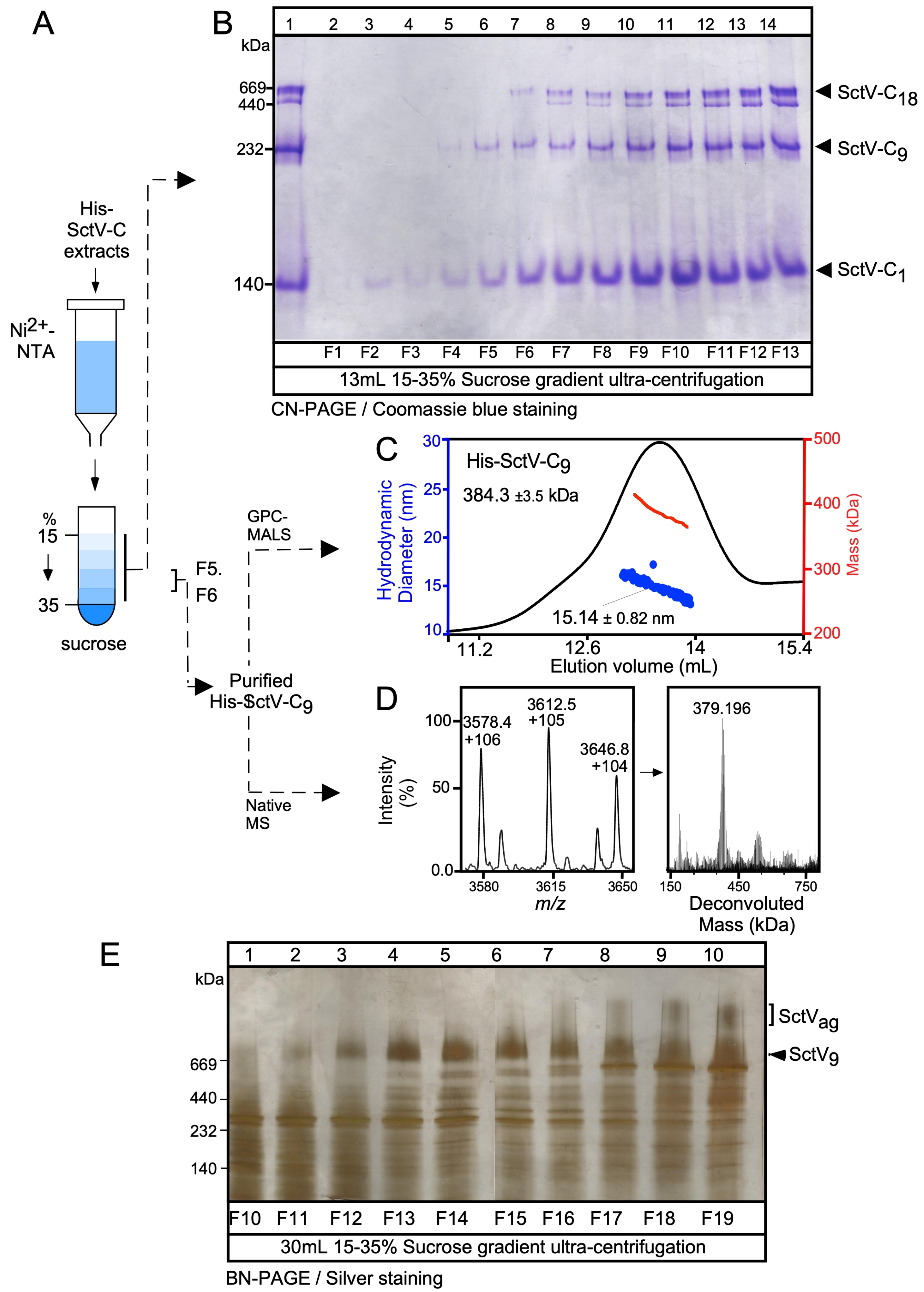
A
His-SctV-C extracts
Ni2+-NTA
%
15
35
sucrose
F5. F6
GPC-MALS
Purified His-SctV-C9
Native MS
B
kDa
669
440
232
140
1 2 3 4 5 6 7 8 9 10 11 12 13 14
F1 F2 F3 F4 F5 F6 F7 F8 F9 F10 F11 F12 F13
13mL 15-35% Sucrose gradient ultra-centrifugation
SctV-C18
SctV-C9
SctV-C1
CN-PAGE / Coomassie blue staining
C
His-SctV-C9
384.3 ±3.5 kDa
15.14 ± 0.82 nm
Hydrodynamic Diameter (nm)
Mass (kDa)
Elution volume (mL)
D
3578.4 +106
3612.5 +105
3646.8 +104
379.196
Intensity (%)
m/z
Deconvoluted Mass (kDa)
E
kDa
669
440
232
140
1 2 3 4 5 6 7 8 9 10
F10 F11 F12 F13 F14 F15 F16 F17 F18 F19
30mL 15-35% Sucrose gradient ultra-centrifugation
SctVag
SctV9
BN-PAGE / Silver staining

**Fig. S4: Purification of $SctV_9$ and SctV-$C_9$ (related to Figures 2, 3 and 4)**

**A**. Purification of the C-terminal cytoplasmic domain His-SctV-$C_9$ using sucrose gradient (15-35% w/v) centrifugation. 500*µ*l (10 mg/mL) His-SctV-C purified from $Ni^{2+}$-NTA resin was loaded on top of a pre-formed sucrose gradient and harvested in 13 fractions (1mL each).

**B**. Fractions F1 to F13 from A, (20*µ*l loaded) were analyzed by CN-PAGE and stained with coomassie blue (see Materials and Methods for details). Fractions with the dimerized 9-meric species were discarded (F6-F13, lanes 7 to 14), while fractions containing nonameric and monomeric species (F5 and F6, lanes 4 and 5) were used for further analysis in C and D. L, loading sample; the SctV-$C_{18}$, SctV-$C_9$, and SctV-C are indicated. The monomeric species migrates aberrantly in CN-PAGE near the position of the 140kDa protein marker but is resolved as a clear monomer by gel permeation chromatography coupled on line with MALS as a species of 45kDa (-/+2). Representative images are shown; *n=5*

**C.** Purified pooled fractions F5/F6 of His-SctV-$C_9$ from A and B (around 50 µM; Buffer M) were analyzed using GPC with online MALS and QELS measurements. SctV-C migrated as a broad peak with a determined mass of 380.9 ± 3.8 kDa and with a hydrodynamic radius of 15.04 ±0.82 nm that would be close to the theoretically expected values of a 9-mer (SctV-$C_9$; theoretical mass 365,34 kDa). UV traces (black) and hydrodynamic radius (blue); Mass (red). *n=3*

**D**. Electrospray Ionization Mass Spectrometry (ESI-MS) analysis of native His-SctV-$C_9$ purified in B. The spectrum of His-SctV-$C_9$ at different charge states (left panel) and its deconvolution into a native mass (right panel) are shown. *n=3*

**E.** Analysis of fractions of full-length His-$SctV_9$ after separation in a sucrose gradient (15-35% w/v) upon ultra-centrifugation. Membrane proteins were extracted (1.8% Triton X-100) from membranes prepared from C41/pACYCDuet-1-*his-sctV* cells after disruption by French press and ultracentrifugation. Detergent-solubilized membrane proteins were separated using a linear sucrose gradient. Sucrose gradient fractions (1mL each; 20*µ*l loaded on the gel) were analyzed by BN-PAGE and stained with silver staining (lanes 1-10; sucrose gradient fractions #10-19). Higher order aggregates of SctV (bracket; $SctV_{ag}$) were observed in the sucrose gradient fractions F16-19 (lanes 7-10). Fractions F10-15 were collected and pooled for subsequent reconstitution into peptidiscs. A representative gel is shown; *n=5.*

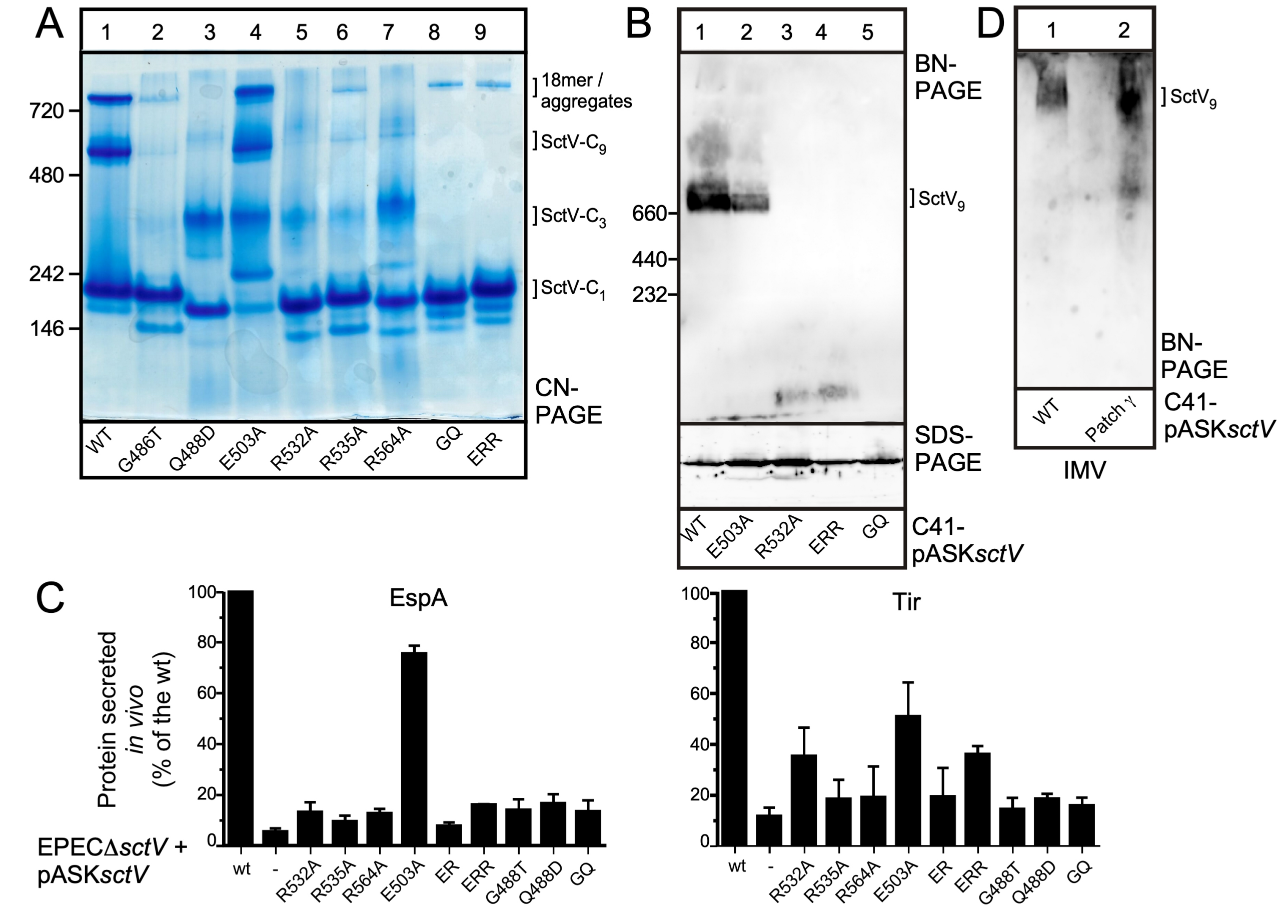


**Fig. S5: Analysis of SctV oligomerization mutants (related to Figure 2).**

**A.** CN-PAGE analysis of His-SctV-C wt or mutated at the oligomerization interface (see Fig. S9 below) synthesized in C41 cells. The protein complexes derived from C41/*sctV-C* wt or mutated were purified using Ni-NTA resin and analyzed by CN-PAGE. With the exception of the E503A derivative (lane 4), all other SctV- C derivatives mutated on the oligomerization interface fail to form higher oligomeric species, and display compromised oligomeric complex formation. ERR: E503A-R535A-R564A; GQ: G486T-Q488D. A representative gel is shown; *n=3*

**B.** BN-PAGE analysis of SctV or of the indicated derivatives mutated in the oligomerization interface synthesized in C41 cells. The membrane fraction of C41 cells expressing plasmid-borne SctV or mutant derivatives was extracted with 1.8% DDM. Samples were analyzed by BN-PAGE (upper panel) and SDS-PAGE (lower panel) and immuno-stained with $\alpha$-SctV-C antibodies. With the exception of the E503A derivative (lane 2) the other SctV-C mutants fail to form higher oligomeric species, abolishing nonameric complex formation of the full-length protein, despite the fact that all derivatives can be produced intracellularly, like the wild type polypeptide. This indicated that C-domain oligomerization is the driving force for SctV oligomerization. ERR: E503A-R535A-R564A; GQ: G486T-Q488D. A representative gel is shown; *n=6*

**C**. Secretion of EspA and Tir from EPECΔ*sctV* cells carrying *sctV* or the indicated mutant derivatives. Secreted protein amounts were quantified and secretion derived from EPECΔsctV/ *sctV* was considered 100%. All other values are expressed as % of this. With the exception of the E503A derivative, the rest of the SctV cytoplasmic domain mutants fail to complement protein secretion through T3SS, suggesting that SctV oligomerization is essential for T3SS function corroborating previous observations. ER: E503A-R535A; ERR: E503A-R535A-R564A; GQ: G486T-Q488D. *n=3*.

**D.** BN-PAGE analysis of SctV or SctV Patch $\gamma$-. C41-IMVs carrying SctV or mutant were treated with 1.8% DDM. Samples were analyzed by BN-PAGE and immuno-stained with $\alpha$-SctV-C antibodies. Mutant SctV can form higher oligomeric species like the wt. A representative gel is shown; *n=5*

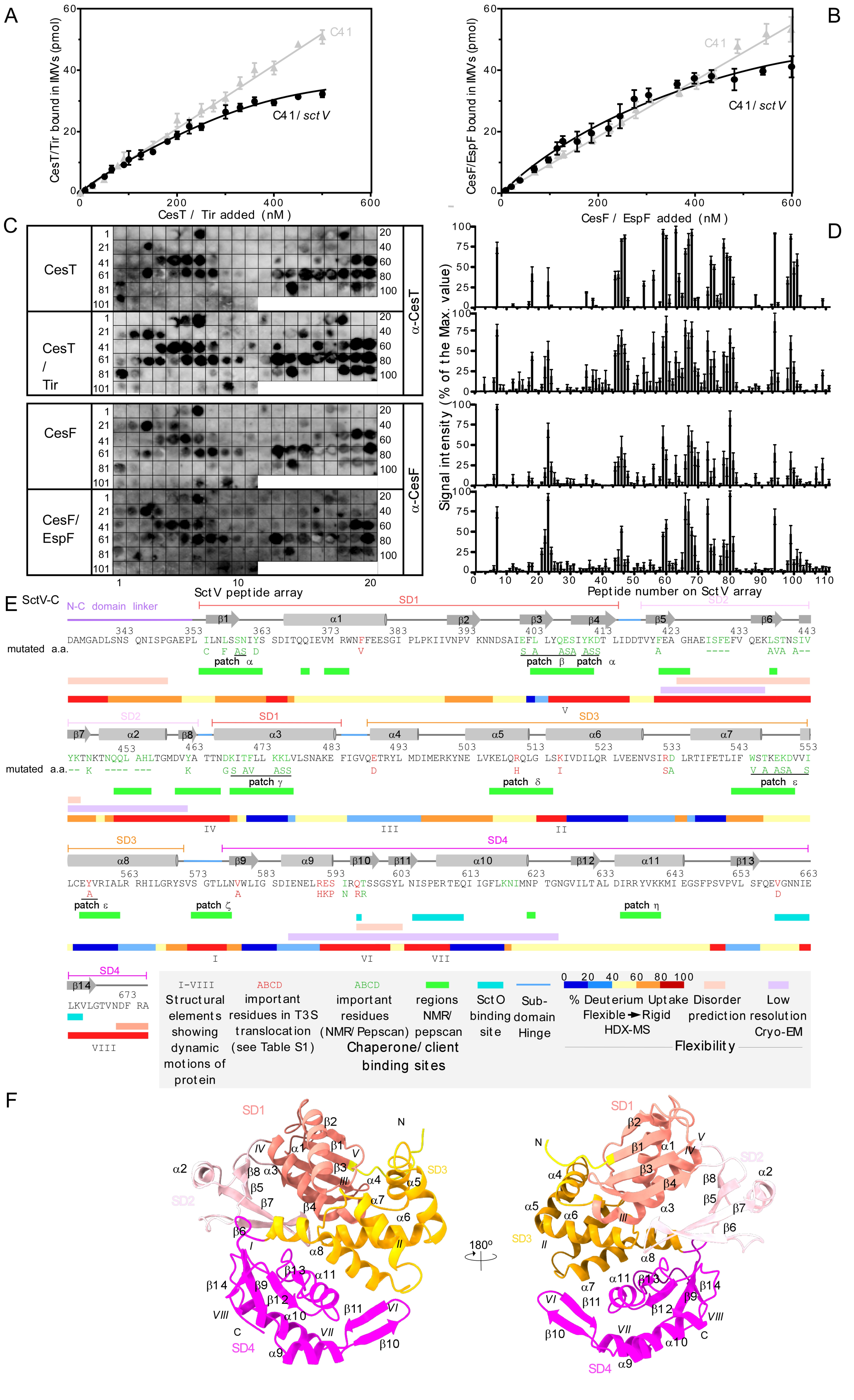

A
CesT/Tir bound in IMVs (pmol)
CesT / Tir added (nM)
C41
C41/ sct V
B
CesF/EspF bound in IMVs (pmol)
CesF / EspF added (nM)
C41
C41/ sct V
C
CesT
CesT / Tir
CesF
CesF/ EspF
α-CesT
α-CesF
SctV peptide array
D
Signal intensity (% of the Max. value)
Peptide number on SctV array
E
SctV-C
N-C domain linker
SD1
SD2
SD3
SD4
mutated a.a.
patch α
patch β
patch γ
patch δ
patch ε
patch ζ
patch η
DAMGADLSNS QNISPGAEPL ILNLSSNIYS SDITQQIEVM RWNFFEESGI PLPKIIVNPV KNNDSAIEFL LYQESIYKDT LIDDTVYFEA GHAEISFEFV QEKLSTNSIV
YKTNKTNQQL AHLTGMDVYA TTNDKITFLL KKLVLSNAKE FIGVQETRYL MDIMERKYNE LVKELQRQLG LSKIVDILQR LVEENVSIRD LRTIFETLIF WSTKEKDVVI
LCEYVRIALR RHILGRYSVS GTLLNVWLIG SDIENELRES IRQTSSGSYL NISPERTEQI IGFLKNIMNP TGNGVILTAL DIRRYVKKMI EGSFPSVPVL SFQEVGNNIE
LKVLGTVNDF RA
I-VIII Structural elements showing dynamic motions of protein
ABCD important residues in T3S translocation (see Table S1)
ABCD important residues (NMR/ Pepscan)
regions NMR/ pepscan
SctO binding site
Sub-domain Hinge
% Deuterium Uptake Flexible → Rigid HDX-MS
Disorder prediction
Low resolution Cryo-EM
Chaperone/ client binding sites
Flexibility
F
SD1
SD2
SD3
SD4
N
C
180°

**Fig. S6: Binding of chaperone/exported client complexes to full-length, membrane-embedded SctV and to an immobilized array of SctV peptides (related to Figures 2, 5 and 6).**

**A** and **B**. Affinity measurements of chaperone/exported client complexes for membrane-embedded full length His-SctV. The binding curves of [$^{35}$S]-CesT/Tir and [$^{35}$S]-CesF/EspF (at concentration ranges of 0-600 nM) for Inner Membrane Vesicles (IMVs) derived from C41 cells with (black) or without (gray) embedded His-SctV$_9$, respectively. Data were analyzed by nonlinear regression in GraphPad Prism 6.0 as described (Portaliou et al., 2017) and represent average values with error bars, standard mean error (SEM) are shown. The interaction of CesT/Tir and CesF/EspF with C41 IMVs exhibited linear, non-saturable binding curves, which did not converge to any $K_d$. *n = 6-9*.

**C-E.** Determination of the chaperone/exported client binding sites on SctV-C using an immobilized peptide array.

**C.** Peptide arrays were used to identify potential chaperone/exported client binding regions on SctV-C. The immobilized peptides (13 mers, with 10 residue overlapping; PepSPOTS™ on Cellulose; JPT) of SctV-C were probed with the indicated purified protein ligands (200 nM; 25 mL Buffer J; 25°C, 1h): CesT, CesT/Tir, CesF, and CesF/EspF, respectively, as described (Portaliou et al., 2017). Proteins bound to the immobilized peptides were electro-transferred onto PVDF membranes (Amersham GE Life sciences) and immuno-stained with $\alpha$-CesT and $\alpha$-CesF antibodies, as indicated. Representative experiments are shown; *n= 6*.

**D.** The intensity of each peptide obtained from **C** was quantified using ImageQuant software (GE Healthcare). All values per experiment were expressed as a percentage of the highest value and plotted using Graph Pad Prism 6.0; Mean values with error bars (SEM) are shown.

**E.** Linear map of C-domain of SctV (residues 334-675) including secondary structure elements (numbered only for the presentation and ignoring the unknown elements of the N-terminus). Chaperone binding regions from pepscans (panel C) and from NMR studies of FlhA-C in flagellum (Xing et al., 2018), after sequence alignment with $_{EPEC}$SctV-C, are shown with green lines below the linear sequence of $_{EPEC}$SctV-C. Similarly, SctO binding regions (Jensen et al., 2020), disordered regions (as predicted by IUpred), flexible, intermediate and rigid areas (derived from HDX-MS analysis; coloured as in Fig. 5A) and low resolution cryo-EM regions (Fig. S8) are shown in different colors, as indicated. In addition, structural elements involved in dynamic motions of proteins are shown in latin numerals (I-VIII). The SctV residues mutated in Patch $\alpha$, $\beta$ and $\gamma$ and the chaperone binding regions defined by NMR are indicated in the primary sequence and below them the substituted residues in the mutant derivatives generated [green; (Portaliou et al., 2017; Xing et al., 2018)]. Additionally, after sequence alignment, we indicate important residues for the gatekeeper interaction (red letters) as identified on SctV homologues [see Table S2 for details; (Lee et al., 2014; Shen and Blocker, 2016; Yu et al., 2018)] and below them the substituted residues are indicated.

**F:** Secondary structural elements and important dynamic motions derived from cryo-EM and HDX-MS analysis respectively are shown in SctV-C protomer. SctV-C sub-domains are colored in a ribbon representation as in Fig. 4E.

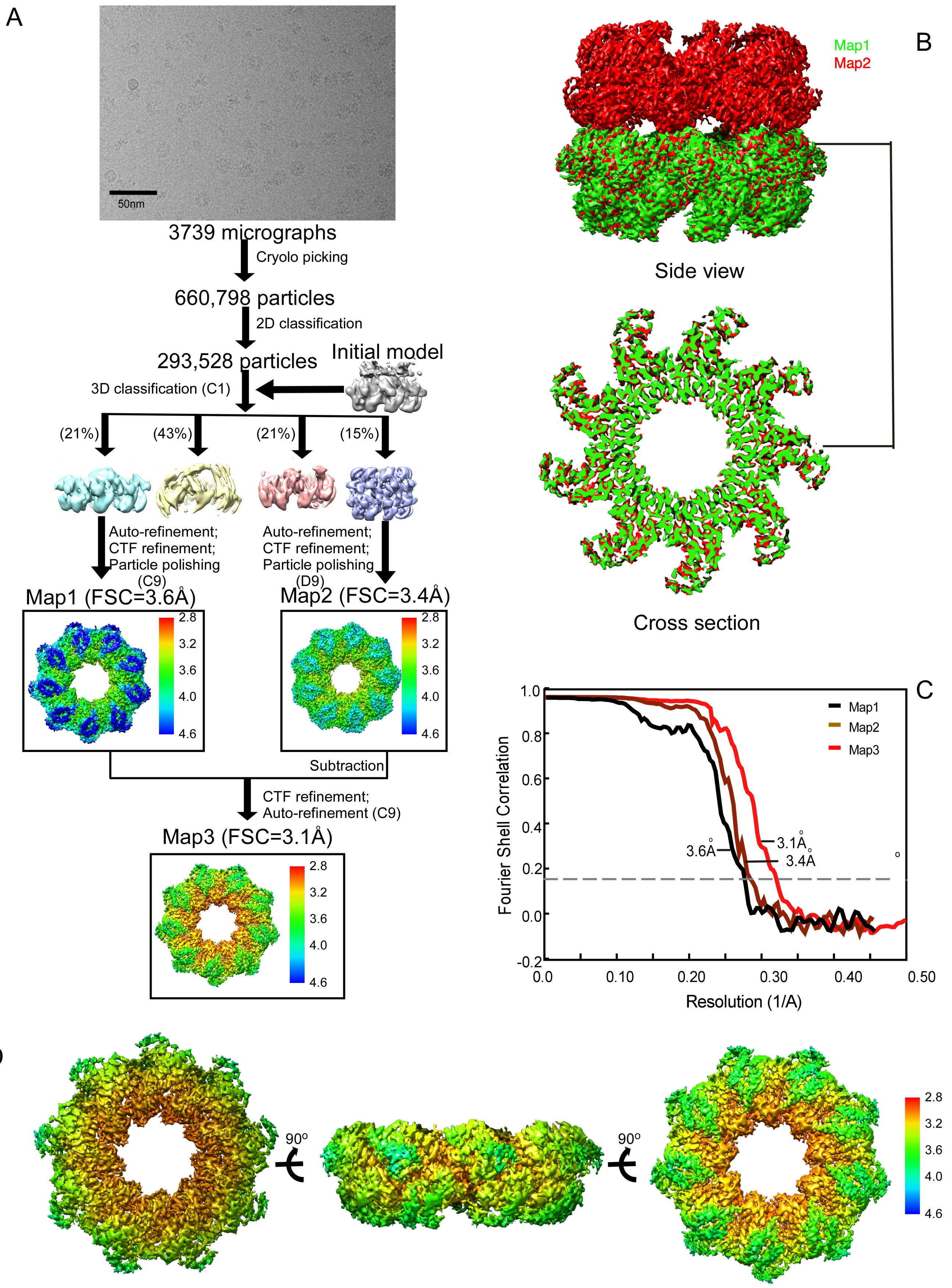


**Fig. S7. Cryo-EM pipeline and structure analysis of SctV-C (related to Figure 4)**

**A.** The cryo-EM data processing workflow.
**B.** Comparison of map1 and map2 reconstructed from single-ringed particles using C9 symmetry and from double ringed particles using D9 symmetry, separately. Cross-section of SctV-C shows that the map1 and map2 are well aligned.
**C.** Gold standard Fourier shell correlation (FSC) curves of the single-ringed map of SctV-C using single-ringed particles (black trace), double-ringed map of SctV-C using double-ringed particles (brown trace), and sing-ringed map of SctV-C using single-ringed particles and subtracted double-ringed particles (red trace).
**D.** Local resolution of the final cryo-EM map (map3) of SctV-C. The map was coloured according to local resolution with the indicated color key. The local resolution was calculated with Relion 3.1 and labeled in ChimeraX1.0.

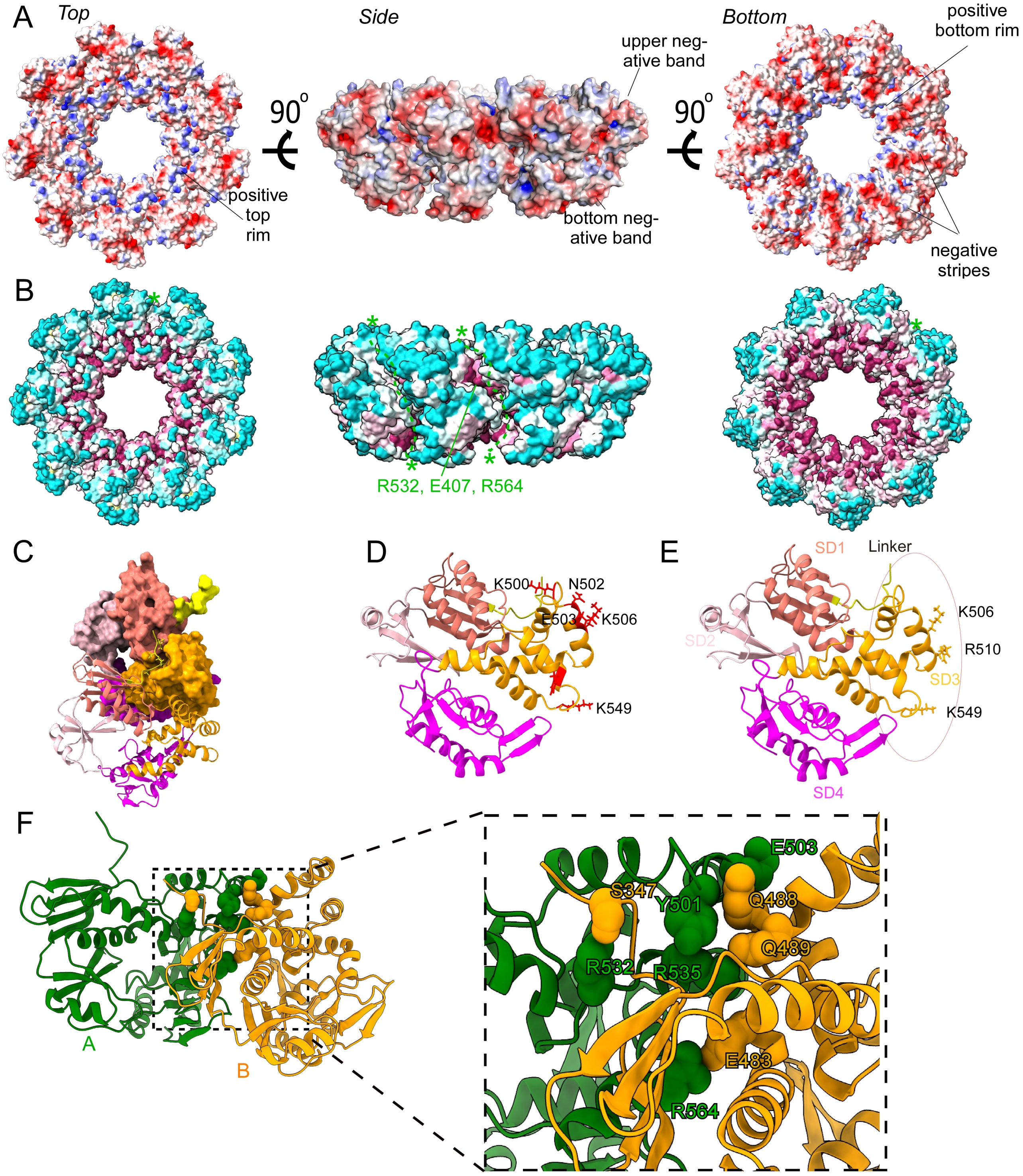


G

| N-C linker-SD3 salt bridge | SD1-SD3 salt bridge | SD3-SD3 salt bridges | SD1-SD3 hinge |
|---|---|---|---|
| S347-R532(A) | E483-R564(A) | Q488(D)-E503(A)/Y501 | 484-487: FIG(T)V |
| | | E489-R535(A) | |

**Fig. S8: Intraprotomeric domain organization/motions in the SctV-C protomer and close-up views of the interproteomeric interaction in the SctV-C nonamer (Related to Figures 2, 4, 6).**

**A.** Electrostatic potential analysis of the SctV-C ring.The surface of the SctV-C ring was coloured by the electrostatic surface potential using ChimeraX. Possitive amino acids are Colored in blue, negative red and neutral white.

**B.** Sequence conservation analysis of the SctV-C ring. ConSurf (https://consurf.tau.ac.il/) was used to estimate the evolutionary conservation of SctV-C. The SctV-C ring is shown as a surface and is colored according to the ConSurf evolutionary conservation score (magenta very conserved to light blue no conservation). R532, E407 and R564 are shown with an asterisk in two adjacent protomers and a dashed line indicates the oligomerization interface.
**C.** Oligomerization interface of two adjacent protomers. The linker region of SctV-C protomer lying inside the small groove formed by SD1 and SD3 of its adjacent protomer. One protomer is shown as surface while the adjacent one in ribbon and both are colored as in Fig. 4B.
**D.** The side chains of the most mobile residues of SctV-C SD3 are shown and coloured in red
**E.** Ribbon diagram indicating the sub-domains of SctV-C (SD; SD1: 353-415 and 463-483; SD2:416-463; SD3: 488-570; SD4: 570-676). The protomer is shown in side view oriented as the orange protomer in 4A, side view**.** Conserved residues of SD3 that their side chains are protruding in the SctV pore are indicated.
**F.** The intermolecular electrostatic interactions that connect a protomer (A; green) to its neighbouring protomer (B; yellow) of EPECSctV-C shown here as spheres in the SctV-C model.
**G**. Mutagenesis analysis of the salt-bridges shown in D and of the flexible SD1-SD3 hinge (484-FIGV-487). The mutated residues are indicated in red. EPECSctV oligomerization mutants failed to restore protein secretion since they can not form oligomers, indicating that oligomerization is essential for T3SS (Fig. 2D-F; Fig. S6).

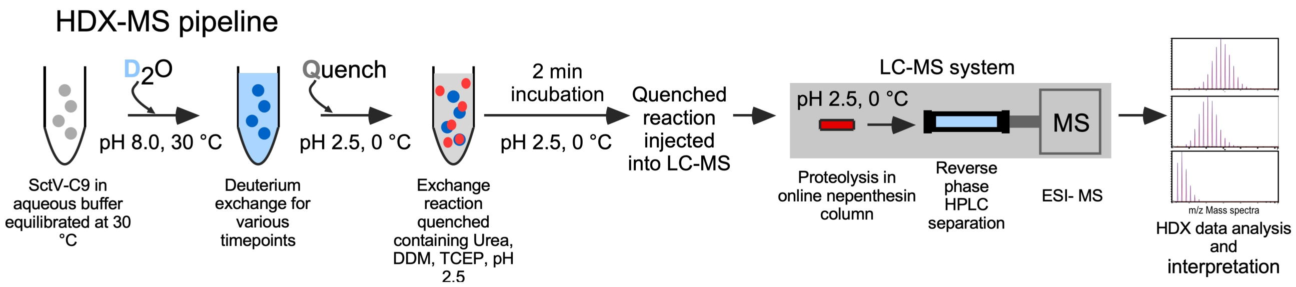


**Fig. S9 HDX-MS pipeline and HDX-MS analysis of *EPEC*SctV-C (related to Figures 4 and 5).**

Schematic representation of the HDX-MS pipeline for analyzing soluble EPECSctV-C. Soluble SctV-$C_9$ (5μM; grey circles) was diluted in deuterated buffer (1:10 dilution for 90% final $D_2O$ concentration; 30°C; final volume 50μL) and incubated for different time points (10 s, 30 s, 1 m, 5 m, 10 m, 30 m, and 1440 m). The reactions were quenched with 50μL Buffer O, incubated for 2 min (4°C) and then injected into a LC-MS system (pH 2.5; 0°C; Waters) and digested using an on-line nepenthesin column (2.1 mm ID x 20 mm length, Cat. No.: AP-PC-004, Affipro) and peptides were separated using reverse phase HPLC C-18 analytical column (130 A°, 1.7 mm, 1 x 100 mm, Waters). HDX-MS data were analyzed and interpreted as described in experimental methods.

### Supplementary Table:

#### Table S1. cryo-EM data and model refinement

| **SctV-C C9 map** | |
|---|---|
| **Data collection and processing** | |
| Electron Microscope | Titan Krios |
| Electron Detector | K3 direct detector |
| Voltage (KV) | 300 |
| Pixel size | 0.55 |
| Electron exposure ($e^-$/ Å$^2$) | 41.25 |
| Defocus range($\mu$m) | 0-4 |
| Symmetry imposed | C9 |
| Number of micrographs | 3739 |
| Initial number of particles image | 293,528 |
| Final number of particles | 105,670 |
| Map resolution (Å) | 3.1 |
| FSC threshold | 0.143 |
| Map sharpening B factor (Å$^2$) | -94.52 |
| Map resolution range (Å) | 3.0-4.2 |
| Initial model used | $_{Sf}$SctV-C(4a5p)/$_{St}$SctV-C(2x49) |
| **Model composition** | |
| Non-hydrogen atoms | |
| Protein residues | 2935 |
| Polypeptide chains | 9 |
| Bonds (Å) | 0.010 |
| Angles(°) | 1.267 |
| **Ramachandran plot** | |
| Favoured (%) | 98.77 |
| Allowed (%) | 100 |
| Outliers (%) | 0 |
| **Validation** | |
| MolProbity score | 0.72 |
| Clash score | 0.67 |
| Rotamer outlier (%) | 0 |

#### Table S2. HDX-MS analysis data

| **Data Set** | **SctV-C$_9$** |
|---|---|
| HDX reaction details | 50 mM Tris, 50 mM KCl, $pH_{read}$ = 8, 30 °C |
| HDX time course (min) | 0.166, 0.5,1.0,5.0,10.0,30.0,100.0 |
| HDX control samples | Maximally-labelled control (WT protein) |
| Back-exchange | 20-45% (peptide composition dependent) Average=28% |
| # of Peptides | 233 |
| Sequence coverage | ~99.7% |
| Average peptide length / Redundancy | 8,70 |
| Replicates (biological or technical) | 2(biological), 3 x 3 (technical), |
| Repeatability (average standard deviation) | 0,05 |
| Significant differences in HDX (delta HDX > X D) | 0.5 D (99.8% CI) |

#### Table S3: SctV residues important for interactors to bind (related to Figures 6 and S6).

| **Mutated aa in SctV** | **Organism** | **Residue in $_{EPEC}$SctV** | **Ref.** | **Phenotype** | |
|---|---|---|---|---|---|
| | | | | **Binding of interactors** | **Secretion** |
| N373D | *S. flexneri* | Y362 | (Shen and Blocker, 2016) | No interaction with SctW | middle substrates secretion: No<br>late substrates secretion: Yes |
| I674V | *S. flexneri* | V658 | | | |
| Q608R | *S. flexneri* | Q596 | | | |
| Q626R | *P. aeruginosa* | | (Lee et al., 2014) | No interaction with SctW domain 1 (Pcr1 protein) | late substrates secretion: Yes |
| G368C | *S. typhimurium* (Flagellum) | I354 | (Inoue et al., 2019; Minamino and Namba, 2008) | Strong interaction with early substrates | Middle substrates secretion: No<br>Late substrates secretion: No |
| D456V | | S438 | (Kinoshita et al., 2013) | Disturbs the binding site of chaperone-exported protein complexes | Reduced secretion and motility |
| F459A | | S441 | | | |
| T490M | | F471 | | | |
| I440A | | F421 | (Xing et al., 2018) | Disturbs the binding site of chaperone-exported protein complexes | Reduced motility |
| F459A | | L437 | | | |
| L461A | | T439 | | | |
| V482K | | Y462 | | | |
| V487G | | D467 | | | |
| D456A | | D533 | | | |
| $^{427}$DDLIE$^{43}$ | *B. subtilis* (Flagellum) | $^{428}$ISFEF$^{432}$ | (Bange et al., 2010) | Disturbs the binding site of chaperone-exported protein complexes | Reduced motility |
| $^{445}$KWISE$^{44}$ | | $^{442}$IVYKT$^{446}$ | | | |
| $^{453}$DEADM$^{4}$ | | $^{450}$NQQLAHL$^{456}$ | | | |
| F378V | *S. typhimurium* | F378 | (Yu et al., 2018) | Disturbs the binding site of gatekeeper SctW | Middle substrates secretion: No<br>Late substrates secretion: Yes<br>(phenocopying secretion profile of gatekeeper deletion) |
| E488D | | E489 | | | |
| R531S | | R532 | | | |
| R509H | | R510 | | | |
| K515I | | K516 | | | |
| R590H | | R591 | | | |
| E591K/ L357F | | E592/L357 | | | |
| S592P | | S593 | | | |
| M450K/ I593N | | N450/I593 | | | |
| T596R | | T597 | | | |
| V632D | | V632 | | | |

### Supplementary materials:

#### Table S4. Buffers

| | |
|---|---|
| **Buffer A** | 50 mM Tris-HCl pH 8.0; 0.5 M NaCl; 5 mM Imidazole; 5% Glycerol |
| **Buffer B** | 50 mM Tris-HCl pH 8.0; 50 mM NaCl; 5 mM Imidazole; 5% Glycerol |
| **Buffer C** | 50 mM Tris-HCl pH 8.0; 50 mM NaCl; 150 mM Imidazole; 5% Glycerol |
| **Buffer D** | 50 mM Tris-HCl pH 8.0; 50 mM NaCl; 10% Glycerol |
| **Buffer E** | 50 mM Tris-HCl pH 8.0; 50 mM NaCl; 50% Glycerol |
| **Buffer F** | 50 mM NaCl; 50 mM Tris-HCl(pH8.0); 2 mM 6-Aminohexanoic acid; 0.5 mM EDTA, 5 mM Imidazole; pH8.0 |
| **Buffer G** | 50 mM NaCl; 50 mM Tris-HCl pH8.0; 0.9% Triton X-100;5 mM Imidazole |
| **Buffer H** | 50 mM NaCl; 50 mM Tris-HCl pH8.0; 5 mM Imidazole |
| **Buffer I** | 50 mM Trizma base pH 8, 6.4 mM KCl, 170 mM NaCl |
| **Buffer J** | 50 mM Trizma base pH 8, 50 mM KCl, 5mM $MgCl_2$, 5 g/L of sucrose, 100 µg/mL BSA, 0.01% v/v Tween-20 |
| **Buffer K** | 50 mM Trizma base pH 8, 2.7 mM KCl, 137 mM NaCl, 0.01% v/v Tween-20 |
| **Buffer L** | 1M NaCl; 50 mM Tris/HCl pH 8.0 |
| **Buffer M** | 50 mM NaCl; 50 mM Tris-HCl pH8.0 |
| **Buffer N** | 50mM Tris-HCl pH 8.0; 50mM KCl; 5mM $MgCl_2$ |
| **Buffer O** | 8MUrea; 5 mM TCEP; 0.1% DDM; Formic Acid 0.7% |
| **1X PBS** | 137 mM $NaCl_2$; 2.7 mM KCl; 4.3 mM $Na_2HPO_4$; 1.4 mM $KH_2PO_4$ |
| **1XM9 salts** | 33.7 mM $Na_2HPO_4$; 22 mM $KH_2PO_4$; 8.55 mM NaCl; 9.35 mM $NH_4Cl$ |
| **Assembly Buffer** | NSPr (an amphipathic bi-helical peptide) mix (1 fluorescent NSPr+ 2 non-fluorescent NSPr) in 20 mM Tris-HCl pH 8.0 |
| **Solubilization buffer** | 50 mM NaCl; 50 mM Tris-HCl pH7.0; 2 mM 6-Aminohexanoic acid; 1mM EDTA; 10% glycerol |
| **BN-PAGE loading buffer** | 5% w/v Coomassie blue G-250 in 500mM 6-Aminohexanoic acid |
| **Cathode buffer 1** | 50 mM Tricine pH 7.0; 7.5 mM Imidazole pH7.0; 0.02% w/v Coomassie blue G-250 |
| **Cathode buffer 2** | 50 mM Tricine pH 7.0; 7.5 mM Imidazole pH7.0; 0.002% w/v Coomassie blue G-250 |
| **Anode buffer** | 25 mM Imidazole pH 7.0 |

#### Table S5. Antisera

Rabbit polyclonal antibodies against the indicated purified proteins or protein domains were raised by Davids Biotechnologie, Germany. Antibodies against T3SS-related proteins were further purified by 9 cycles of negative immuno-absorption, using membranes isolated from EPEC strains that lacked the gene of interest, i.e. for $\alpha$-SctJ, membranes isolated from EPECΔ*sctJ* cells were used.

| Antibody | Animal Source | Reference or commercial source |
|---|---|---|
| α-SctV-C | Rabbit | (Portaliou et al., 2017) |
| α-SctU-C | Rabbit | This study |
| α-GFP | Rabbit | (Hamed et al., 2018) |
| α-CesT | Rabbit | (Portaliou et al., 2017) |
| α-CesF | Rabbit | (Portaliou et al., 2017) |
| α-Rabbit IgG | Goat | Jackson ImmunoResearch Europe Ltd |
| α-His tag | Mouse | SEROTEC |
| α-Mouse IgG | Goat | Jackson ImmunoResearch Europe Ltd |

#### Table S6. Bacterial strains

| Bacterial strain *E. coli* | Description (gene deleted) | Reference/source |
|---|---|---|
| DH5a | F$^-$ *endA1 glnV44 thi-1 recA1 relA1 gyrA96 deoR nupG purB20* φ80d*lacZ*ΔM15 Δ(*lacZYA-argF*)U169, hsdR17($r_K^- m_K^+$), λ$^-$ | Invitrogen |
| BL21(DE3) | *E. coli* str. B F$^-$ *ompT gal dcm lon hsdS$_B$*($r_B^- m_B^-$) λ(DE3 [*lacI lacUV5-T7p07 ind1 sam7 nin5*]) [*malB*$^+$]$_{K-12}$(λ$^S$) | (Studier et al., 1990) |
| C41(DE3) | F$^-$ *ompT gal dcm hsdS$_B$*($r_B^-$ $m_B^-$)(DE3) | Lucigen (Miroux and Walker, 1996) |
| EPEC E2348/69 | *E. coli* O127:H6 (strain E2348/69) | (Levine et al., 1978) |
| EPEC E2348/69 Δ*sctV* | Δ*sctV*:: *nptII*(Kan$^R$) | (Portaliou et al., 2017) |

**Table S7. Vectors and genetic constructs:**

Genes were cloned in the indicated plasmid vectors using a combination of restriction sites (as indicated). DNA restriction enzymes and DNA polymerases were from Promega. All PCR-generated plasmids were sequenced (Macrogen, Europe). Plasmids were transformed in DH5α cells and stored in 20% glycerol at –80°C.

| Vector name | Antibiotic resistance | Promoter | Origin | Reference/source |
|---|---|---|---|---|
| pASKIBA7 | Amp | Tet | pBR322 | IBA life sciences; (Guzman et al., 1995) |
| pETDuet-1 | Amp | T7 | pBR322 | Novagen |
| pACYCDuet-1 | Cm | T7 | p15A | Novagen |
| pET16b | Amp | T7 | pBR322 | Novagen |
| pBAD501 | Gm | pBAD | p15A | (Chatzi et al., 2017) |
| pBAD24 | Amp | pBAD | pBR322 | (Guzman et al., 1995) |
| **Gene** | **Uniprot KB accession** | **Plasmid name** | **Vector** | **Cloning strategy or source** |
| *sctV* | B7UMA7 | pLMB1823 | pACYCDuet-1 | The *sctV* gene was digested out from pETDuet1-*sctV* (pLMB0057) and inserted in pACYCDuet1 after NdeI-XhoI digestion. |
| *sctV* | B7UMA7 | pLMB0088 | pASKIBA 7 | (Portaliou et al., 2017) |
| *sctV* $\gamma-$ | B7UMA7 | pLMB1764 | pASKIBA 7 | (Portaliou et al., 2017) |
| *sctV* $\varepsilon-$ | B7UMA7 | pLMB1765 | pASKIBA 7 | The synthetic gene fragment of *sctV* (sgLMB017) was digested out and inserted to pASK sctV- no HindIII (pLMB1912). |
| *his-sctV* | B7UMA7 | pLMB1840 | pACYCDuet-1 | The *sctV* gene was amplified from EPEC E2348/69 (B1030) using primers X2126 and X1710. The gene was inserted at the 2nd multiple cloning site of pACYCDuet-1 after NdeI and XhoI digestion. |
| *his-sctV* | B7UMA7 | pLMB1877 | pASKIBA500 | The *his-sctV* gene was digested out from pLMB1840 (pACYCDuet1-*his-sctV*) using NdeI-XhoI restriction enzymes and inserted in pASK500 at the same sites. |
| *his-sctV* | B7UMA7 | pLMB1841 | pBAD501 | The *his-sctV* gene was digested out from pLMB1840 (pACYCDuet1-*his-sctV*) using NdeI-XhoI restriction enzymes and inserted in pBAD501 at the same sites. |
| *his-sctRSTU* | B7UMB8<br>B7UMB9<br>B7UMC0<br>B7UMC1 | pLMB1824 | pET Duet 1 | The *sctR/S/T/U* genes were amplified from EPEC E2348/69 (B1030) using primers X1725 and X1527. The genes were inserted |

| | | | | |
|---|---|---|---|---|
| | | | | at the 2nd multiple cloning site of pet Duet-1 after NcoI and BamHI digestion. |
| *his-sctV*(N1-333) | B7UMA7 | pLMB1881 | pET16b | The *sctV* (N1-333) fragment was amplified from EPEC E2348/69 (B1030) using primers X1709 and X2132. The gene was inserted in pET16b after NdeI and XhoI digestion. |
| *his-sctVc*(N334-675) | B7UMA7 | pLMB1676 | pET16b | (Portaliou et al., 2017) |
| *his-sctVc* (E503A) | B7UMA7 | pLMB2140 | pET16b | pLMB1676 was used to generate the *his-sctVc(*E503A*)* mutant by site directed mutagenesis |
| *his-sctVc* (R532A) | B7UMA7 | pLMB2141 | pET16b | pLMB1676 was used to generate the *his-sctVc(*R532A*)* mutant by site directed mutagenesis |
| *his-sctVc* (R535*A)* | B7UMA7 | pLMB2142 | pET16b | pLMB1676 was used to generate the *his-sctVc(*R535A*)* mutant by site directed mutagenesis |
| *his-sctVc* (R564A*)* | B7UMA7 | pLMB2143 | pET16b | pLMB1676 was used to generate the *his-sctVc(R564A)* mutant by site directed mutagenesis |
| *his-sctVc* (E503A/R535A*)* | B7UMA7 | pLMB2144 | pET16b | pLMB2140 was used to generate the *his-sctVc(*E503A/R535A) mutant by site directed mutagenesis |
| *his-sctVc* (E503A/R535A/R564A*)* | B7UMA7 | pLMB2145 | pET16b | pLMB2144 was used to generate the *his-sctVc(*E503AR/535AR/564A) mutant by site directed mutagenesis |
| *his-sctVc* (G486T) | B7UMA7 | pLMB2137 | pET16b | pLMB1676 was used to generate the *his-sctVc(*G486T*)* mutant by site directed mutagenesis |
| *his-sctVc* (Q488D*)* | B7UMA7 | pLMB2138 | pET16b | pLMB1676 was used to generate the *his-sctVc(*Q488D*)* mutant by site directed mutagenesis |
| *his-sctVc* (G486T/Q488D*)* | B7UMA7 | pLMB2139 | pET16b | pLMB2137 was used to generate the *his-sctVc(*G486T/Q488D*)* mutant by site directed mutagenesis |
| *sctV(*E503A*)* | B7UMA7 | pLMB2131 | pASKIBA7 | pLMB0088 was used to generate the *his-sctV(*E503A*)* mutant by site directed mutagenesis |
| *sctV(*R532A*)* | B7UMA7 | pLMB2132 | pASKIBA7 | pLMB0088 was used to generate the *his-sctV(*R532A*)* mutant by site directed mutagenesis |
| *sctV*(R535A*)* | B7UMA7 | pLMB2133 | pASKIBA7 | pLMB0088 was used to generate the *his-sctV(*R535A) mutant by site directed mutagenesis |
| *sctV(*R564A*)* | B7UMA7 | pLMB2134 | pASKIBA7 | pLMB0088 was used to generate the *his-sctV(*R564A) mutant by site directed mutagenesis |
| *sctV(*E503A/R535A) | B7UMA7 | pLMB2135 | pASKIBA7 | pLMB2131 was used to generate the *his-sctV(*E503A/R535A*)* mutant by site directed mutagenesis |
| *sctV(*E503A/R535AR/564A*)* | B7UMA7 | pLMB2136 | pASKIBA7 | pLMB2135 was used to generate the *his-sctV(*E503A/R535A/R564A*)* mutant by site directed mutagenesis |
| *sctV(*G486T*)* | B7UMA7 | pLMB2128 | pASKIBA7 | pLMB0088 was used to generate the *his-sctV(*G486T*)* mutant by site directed mutagenesis |
| *sctV(*Q488D) | B7UMA7 | pLMB2129 | pASKIBA7 | pLMB0088 was used to generate the *his-sctV(*Q488D*)* mutant by site directed mutagenesis |

| | | | | |
|---|---|---|---|---|
| *sctV(*G486T/Q488D) | B7UMA7 | pLMB2130 | pASKIBA7 | pLMB0088 was used to generate the *his-sctV(*G486T/Q488D*)* mutant by site directed mutagenesis |
| *eyfp-sctV* | B7UMA7 | pLMB1800 | pASK IBA500-*eyfp*-MCS | The *sctV* gene was digested out from pLMB0088 (pASKIBA7-*sctV*) and was inserted in pLMB1774 (pASK500-*eyfp*-MCS) after NheI-XhoI digestion. |
| *torA-gfp* | --- | --- | pBAD24 | (Barrett et al., 2003) |
| *lacY-eyfp* | --- | --- | pBAD/His | (Robinson et al., 2015) |
| *eyfp* | --- | pLMB1774 | pASKIBA500 | The 800bp *eyfp* gene was amplified from pLMB1707 using primers X2058 and X2062. The gene was inserted in pASKIBA500 (pLMB0014) after XbaI-NdeI digestion. |
| *his-cesT* | P21244 | pIMBB1157 | pETDuet-1 | (Portaliou et al., 2017) |
| *his-cesT and tir* | P21244 and B7UM99 | pIMBB1158 | pETDuet-1 | (Portaliou et al., 2017) |
| *bamE-gfp* | P0A937 | pDR*bamE*-GFP | pDRGFP | R. Ieva |
| *his-cesF* | B7UM99 | pIMBB664 | pETDuet-1 | (Portaliou et al., 2017) |
| *his-espF* | B7UM88 | pIMBB612 | pET22b | The *espF-his* gene was amplified from EPEC E2348/69 (B1030) using primers X342 and X344 and inserted in pET22b vector after NdeI- XhoI digestion. |

#### Table S8. List of primers used for gene cloning

| **Primer Name** | **Gene** | **Restriction site** | **Forward/ Reverse** | **Sequence (5'-3') (restriction sites underlined/ linker italics/ mutation bold)** |
|---|---|---|---|---|
| X2126 | his-*sctV* | *NdeI* | Forward | GGAATTCCATATGCATCACCATCATCACCACATGAATAAACTCTTAAATATATTTAAAA |
| X1710 | *his-sctV* | *XhoI* | Reverse | GGCCTCGAGTCATGCTCTGAAATCATTTAC |
| X1709 | *his-sctV*(N1-333) | *NdeI* | Forward | GGAATTCCATATGAATAAACTCTTAAATATATTTAAAA |
| X2132 | *his-sctV*(N1-333) | *XhoI* | Reverse | GACCCGCTCGAGTTACTTATTATTGCCAGCTCCAAAT |
| X342 | *espF* | *NdeI* | Forward | GGGAATTCCATATGCTTAATGGAATTAGTAACGCTG |
| X344 | *espF* | *XhoI* | Reverse | CCGCTCGAGCCCTTTCTTCGATTGCTCATAGGC |
| X2058 | *eyfp* | *XbaI* | Forward | GCGGTCTAGATAACGAGGGCAAAAAATGGGTAGCATGGT |
| X2062 | *eyfp* | *NdeI* | Reverse | GGGAATTCCATATG*AGAGCCTCCGCCAGAGCCTCCGCC*CTTGTACAGCTCGTCCATGCCGAG |
| X1725 | *his-sctRSTU* | *NcoI* | Forward | CATGCCATGGGCAGCAGCCATCACCATCATCACCACATGTCTCAATTAATGACCATTGG |
| X1527 | *his-sctRSTU* | *BamHI* | Reverse | CGCGGATCCTTAATAATCAAGGTCTATCGCAATAC |
| - | *sctVE503A* | | Forward | GAGAGAAAATATAAC**GCT**CTTGTGAAAGAGCTG |
| - | *sctVE503A* | | Reverse | CAGCTCTTTCACAAG**AGC**GTTATATTTTCTCTC |
| - | *sctVR532A* | | Forward | GAAAATGTCTCAATT**GCA**GATCTGAGAACTATC |
| - | *sctVR532A* | | Reverse | GATAGTTCTCAGATC**TGC**AATTGAGACATTTTC |
| - | *sctVR535A* | | Forward | TCAATTAGAGATCTG**GCA**ACTATCTTTGAGACG |
| - | *sctVR535A* | | Reverse | TCAATTAGAGATCTG**GCA**ACTATCTTTGAGACG |
| - | *sctVR564A* | | Forward | CGTATCGCCCTGCGT**GCT**CATATTTTAGGTCGC |
| - | *sctVR564A* | | Reverse | GCGACCTAAAATATG**AGC**ACGCAGGGCGATACG |
| - | *sctVG486T* | | Forward | GCCAAAGAGTTCATC**ACT**GTACAAGAAACGCGT |
| - | *sctVG486T* | | Reverse | ACGCGTTTCTTGTAC**AGT**GATGAACTCTTTGGC |
| - | *sctVQ488D* | | Forward | GAGTTCATCGGCGTA**GAT**GAAACGCGTTATTTG |
| - | *sctVQ488D* | | Reverse | CAAATAACGCGTTTC**ATC**TACGCCGATGAACTC |
| - | *sctVG486TQ488D* | | Forward | GAGTTCATCACTGTA**GAT**GAAACGCGTTATTTG |

| | | | | |
|---|---|---|---|---|
| - | *sctVG486TQ488D* | | Reverse | CAAATAACGCGTTTC**ATC**TACAGTGATGAACTC |

#### Supplementary movies:

##### Movie S1: Sub-domain motions of SctV-C

$_{EPEC}$SctV-C was compared to the virulence T3SS homologues $_{Cp}$SctV-C from *Chlamydia pneumoniae* (Jensen et al., 2020)(PDB:6WA6) represennting the "open" state and $_{Sf}$SctV-C(PDB: 2x49) from *Salmonella typhimurium* (Worrall et al., 2010) representing the "closed" state. The structure of $_{EPEC}$SctV-C resolved here by cryo-EM (Fig. 4A) was classified as a "intermediate" state.

##### Movie S2: Highly dynamic regions obtained from HDX-MS analysis.

Only the highly flexible regions derived from the HDX-MS analysis are included (Fig. S6E, red)

##### Movie S3: Chaperone binding sites mapped on one protomer of SctV$_9$

Binding sites are from NMR data (Xing et al., 2018) and patches $\alpha$,$\beta$ and $\gamma$ from pepscan analysis [Fig. S6E; (Portaliou et al., 2017)].

##### Movie S4: Binding sites of the ATPase inner stalk protein SctO mapped on two adjacent protomers of SctV$_9$

### Supplementary methods:

#### Preparation of cells for live-cell fluorescence microscopy

Bacterial strains were transformed with indicated plasmids and grown overnight at 37ºC on LB agar plates supplemented with the appropriate antibiotics. An overnight pre-culture starting from one single colony from the LB plates was inoculated in EZ rich defined medium (1:100 dilution; Teknova) with 0.2% w/v glucose as the carbon source and specific antibiotics until $OD_{600}$ reached 0.3-0.4. Gene expression was induced with AHT (as indicated) at 37ºC for 30-60 min. Cells were pelleted by centrifugation at 3,000 x g for 3 min to remove the inducer and resuspended in fresh EZ rich medium before imaging. Coverslips were firstly cleaned by sonicating in 5M KOH for 1h, and then rinsed with deionized water several times and dried with nitrogen. Following the cleaning step, coverslips were treated by Oxygen plasma cleaning for 10 min (PE-50 Compact Bench-top Plasma Cleaning System, Plasma Etch) to remove contaminants from the glass surfaces. Cells were sandwiched between a clean glass cover slip and an agarose slab (10 mg/mL Agarose; low gelling temperature; Sigma-Aldrich) and imaged with live-cell fluorescence microscopy.

#### Western blotting analysis

Following SDS-PAGE or Native-PAGE analysis, proteins were transferred onto nitrocellulose membrane (PROTRAN) or PVDF membrane (Thermo Scientific) using Semi-Dry Transfer apparatus (BIORAD) following manufacturer's instruction (20V; 20 min for SDS-PAGE; 40 min for Native-PAGE). After antibody staining, membranes were incubated with SuperSignal$^{TM}$ West Pico PLUS chemiluminescent substrate (Thermo Scientific) for 2 min for signal development. Images were acquired using Las4000 (GE Healthcare). For image acquisition the manufacturer's setting were used (Resolution / Sensitivity: standard mode; Exposure time 10-20 min; Image dimensions 210 x 140 mm; Image resolution 176 dpi).

#### Sub-cellular protein localization of SctV cytoplasmic or transmembrane domain

C41 cells carrying either pETDuet-*sctV,* or His-*sctV*N1-333 or His *sctV*-C were induced with 0.2 mM IPTG for 12 hours, at 18ºC. Bacteria were resuspended in Buffer A and lysed with French press. Unbroken cells were removed (3,000 x g; 10 min; 4ºC) before high-speed centrifugation (100,000 x g; 30 min; 4 ºC;45 Ti rotor; Optima XPN-80; Beckman Coulter). Sequential washes with 8M urea overnight and 2 hours at room temperature were introduced, when indicated, to remove the inclusion bodies of expressed proteins associated with the membrane. Equal amounts of cytosolic or membrane proteins were analyzed on 12% acrylamide gels by SDS-PAGE and immuno-stained.

#### Peptide array analysis

Peptide array analysis was performed as described previously (Karamanou et al., 2008; Portaliou et al., 2017). In brief, 13mer-, overlapping peptides of SctV-C were immobilized on cellulose membranes (PepSpot Peptides; JPT Peptide Technologies; Germany). Before screening, the dry membrane was washed in methanol (10 min), in high-salt buffer I at room temperature (3 × 5 min), in buffer I at the desired temperature (5 × 5 min) and finally in equilibration buffer J (15 min). 200 nM of chaperones or the chaperone/exported protein complexes were incubated with the peptide array membrane for 30 min at 25°C. Unbound complexes were removed with extended washes with buffer K. The proteins bound to peptides were electro-transferred onto PVDF membrane (Thermo Scientific). Bound proteins to specific peptides were visualized by western-blot using specific antibodies.

**Determination of equilibrium dissociation constants ($K_d$)**

Equilibrium dissociation constants determination was conducted as described previously (Gouridis et al., 2009; Portaliou et al., 2017). In brief, the [$35^S$]-labelled proteins were labelled *in vitro* using the Easy TagTM L-[$35^S$]-methionine (1 mC, Perkin Elmer) and the TNT® Quick coupled Transcription/Translation systems (Promega), according to the manufacturer's instructions.

Proteins and protein complexes stored in Buffer E were serially diluted in Buffer N (20 concentration points; 0.01-1 µM) and mixed with IMVs (20 µg total membrane protein/reaction; 20 µl in Buffer N). 1µl of [$^{35}S$]-labelled protein was added to all the reactions, as a tracer. Samples were incubated on ice for 20 min, and then overlaid on an equal volume of BSA/sucrose cushion (0.2 M Sucrose; 1 mg/mL BSA in Buffer N) and centrifuged (300,000 x g; 20 min; 4°C; rotor TLA-100; Optima Max-XP, Beckman-Coulter). The pellet (containing IMVs and IMVs-bound proteins) was resuspended in 300 µl Buffer N by using a water-bath sonicator and proteins were immobilized on a nitrocellulose membrane using a vacuum manifold (Bio-Dot apparatus; Bio-Rad). Binding of [$35^S$]-labelled proteins on IMVs was visualized by using a high-resolution phosphor storage screen (GE Healthcare) on a Typhoon FLA 9500 system (GE Healthcare; default settings). Image Quant software (GE Healthcare) was used for signal quantification. Data were analyzed by non-linear regression fit for one binding site, using Prism 6.0 (GraphPad). For the determination of each $K_d$, $n=6$, each using 20 concentration points.

**Native mass spectrometry**

His-SctV-C9 was dialyzed in 50 mM ammonium acetate and analyzed using electrospray ionization mass spectrometer (ESI-MS) with a Q-TOF mass analyzer (Synapt G2 HDMS, Waters) and sodium iodide (2 mg/mL) as a calibrant. The operating parameters for the spectral acquisition were as follows: capillary voltage, 1.8 kV; sample cone voltage, 60 V; extraction cone voltage, 2 V; source temperature, 80°C; desolvation temperature, 150°C; backing pressure, 5.26 mbar; source pressure, 2.08 e-3 mbar; Trap, 1.44 e-3 mbar. Spectra were acquired in the range of 900–8000 m/z in positive-ion V mode. The molecular mass of the recorded spectra was calculated using MassLynx software (MassLynx version 4.1) by deconvoluting the mass spectra using MaxEnt 1.
